# Ecology and sexual conflict drive the macroevolutionary dynamics of female-limited colour polymorphisms

**DOI:** 10.1101/2025.02.15.637939

**Authors:** Beatriz Willink, Tammy Ai Tian Ho, Erik I Svensson

## Abstract

Sexual conflict over mating has been documented in many species, both in the field and in experimental studies. In pond damselflies (family Coenagrionidae), sexual conflict maintains female-limited colour polymorphisms, with one female morph typically being a male mimic. However, it is not known whether sexual conflict can also explain the evolutionary origin of novel female morphs, and if so, what ecological factors play a role in this macroevolutionary transition, by modulating the strength of the conflict. Furthermore, the effects of sexual conflict on phylogenetic diversification remain controversial, with studies arguing that the evolution of alternative reproductive morphs, such as female-limited colour morphs, could either hinder or accelerate speciation. Here, we use phylogenetic comparative methods to show that female colour polymorphisms are more likely to evolve when population densities at breeding sites are high, and that these demographic conditions are more common at high latitudes and in open landscapes. We show that female-limited polymorphisms typically evolve from sexually dimorphic ancestors through the addition of a male-like female morph, consistent with the hypothesis of selection for male mimicry. Female polymorphic and sexually dimorphic lineages diversify at a higher rate than sexually monomorphic lineages, suggesting a broad diversity-promoting role of inter-sexual interactions based on visual signals. We conclude that female colour polymorphisms evolve in a predictable fashion, and are likely driven by ecological conditions that increase the rate of pre-mating interactions and thus the intensity of sexual conflict.

## Introduction

A fundamental goal in evolutionary biology is to understand the link between the processes that fuel microevolutionary change and the patterns of trait variation that occur across groups of related organisms (1–3). One way to bridge this gap is to start from a well-understood microevolutionary force, and ask what phenotypic differences between species should we expect, had this process been causal throughout the evolutionary history of a particular clade (e.g. 4, 5, 6). Through this approach, phylogenetic comparative methods, which have traditionally been viewed as exclusively correlational, can be incorporated into an explicit causal framework (7, 8). Here, we use this causal approach to investigate the macroevolutionary dynamics of female-limited colour polymorphisms in damselflies. Specifically, we asked if the macroevolutionary origins of the polymorphisms could be explained by the same process of sexual conflict over mating rates that maintains sympatric morphs within populations.

Inter-locus sexual conflict arises whenever the fitness consequences of mating and reproduction ultimately differ between the sexes (9). If so, the evolution of traits involved in pre-mating interactions, such as mating harassment, will reflect genetic conflicts of interest between alleles segregating at different loci (10). For example, sexual conflict can drive evolutionary change in male-specific behaviours that bias the outcome of pre-mating interactions in favour of male fitness, and at the expense of female fitness. This will in turn impose selection on other loci, governing female behavioural, morphological, or physiological defence traits against such antagonistic pre-mating interactions (11). One possible outcome of inter-locus sexual conflict is the continuous reiteration of this coevolutionary process, propelling an “arms race” between the sexes (12–16). Alternatively, inter-locus sexual conflict can, under certain circumstances, result in female diversification into multiple genetic clusters, or morphs, to which all males cannot simultaneously adapt (16, 17). Consistent with theory (16, 17), the sympatric co-occurrence of multiple heritable female morphs that differ in mating-related traits has been reported in several different taxa, particularly in birds and insects (18–23).

In damselflies, female-limited colour morphs display alternative reproductive tactics (20), involving a suite of correlated behavioural, morphological, physiological, and life-history traits (24–27). Female-limited colour polymorphisms in damselflies typically include a male-coloured morph and one or more female morphs markedly different from males (28). In the species with microevolutionary field studies, empirical evidence indicates that negative frequency-dependent selection via pre-mating male harassment maintains these genetic polymorphisms (29–33). Visually-oriented mate-searching males are thought to disproportionally direct their mating attempts towards common female morphs, and the excessive mating and pre-mating interactions encountered by targeted females can reduce their fecundity ((31); (34); (35); (36); (37)). If avoidance of male mating harassment is the primary driver of the evolution of female-limited colour polymorphisms in damselflies, then these polymorphisms should generally evolve via the origin of male-like females, that can obtain a frequency-dependent fitness advantage from male mimicry. Moreover, we expect that damselfly lineages with marked male pre-mating harassment would more often evolve and maintain inter-sexual mimicry and thus female-limited colour polymorphisms, compared to lineages with lower intensity of sexual conflict.

Mating interactions are shaped by demographic conditions, particularly population densities and sex ratios (38–42), which can in turn be governed by the physical environment (43–45). Empirical evidence from microevolutionary studies suggests that high population densities and male-biased sex ratios increase the intensity of sexual conflict in several taxa, including damselflies (20, 31, 46–50). Such demographic conditions are expected when reproduction is restricted to a short seasonal window, leading to increased adult synchrony and a greater potential for male-male scramble competition over mating opportunities (51–53). We therefore hypothesized that the concurrence of mating interactions in the warmer months of the year should ultimately drive higher evolutionary rates of female-limited polymorphisms in temperate areas, where breeding seasons are shorter compared to the tropics (Fig. 1a). Apart from temporal constraints on mating opportunities, spatial constraints can also promote sexual conflict over mating rates. For instance, in damselfly species that breed in open habitats, which tend to be relatively homogeneous environments, individuals typically stay at or near their reproductive sites throughout most of their adult life (e.g. 54), resulting in high adult densities and high encounter rates between the sexes (Fig. 1a). Conversely, sexual conflict should become more relaxed in structurally complex environments, where females can more readily escape male mating harassment, such as in forests or other habitats with pronounced three-dimensional structure (55–57).

**Figure 1.**
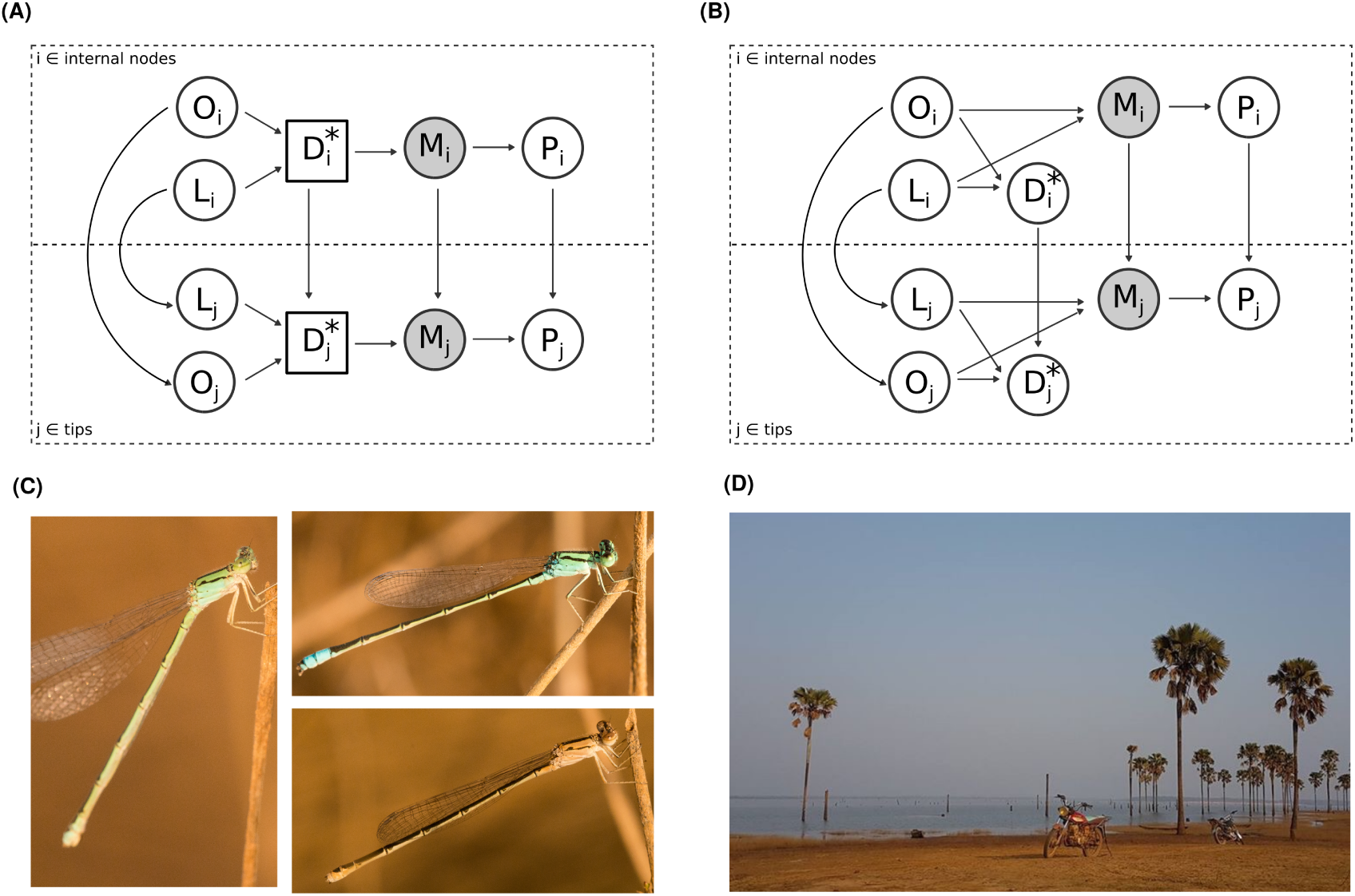
Ecological and demographic factors hypothesized to promote the evolution of female-limited colour polymorphisms in damselflies. **(A, B)** Directed acyclical graphs (DAGs) showing alternative causal scenarios for the role of ecological variation on the evolution of female-limited polymorphisms. Empty circles represent observed variables, and gray circles represent latent variables (not directly quantified). L = latitudinal range, O = habitat openness, D = density at breeding sites, M = male mating harassment, P = presence of female-limited polymorphisms. *The diagrams show density (D) as a demographic predictor of female polymorphism, but a similar causal hypothesis was also addressed using operational sex ratio (OSR) instead. In each scenario, the upper panel represents effects in ancestral lineages (i) and the lower panel shows causal relationships in extant taxa (j). Arrows connecting the two panels thus reflect the role of history (i.e. phylogenetic non-independence). These confounding effects were adjusted by fitting the phylogenetic variance–covariance matrix as a random effect, in Bayesian Phylogenetic Mixed Models (BPMM). **(A)** Density (D) is a mediator (represented by a square node) of the effects of latitudinal range (L) and habitat openness (O) on male-mating harassment (M). **(B)** Latitudinal range (L) and habitat openness (O) impact both density (D) and male-mating harassment (M), but the effects on M are not mediated by D. **(C)** *Pseudagrion nubicum*, an example of a tropical species treated as female-polymorphic in this study. This taxon lacked previous literature data on female-limited colour variation. We sampled 81 individuals of *P. nubicum* at their breeding habitat in Cameroon **(D)** and recorded males (top right), male-like females (left) and non-mimicking females (bottom right). Male-like females were relatively common; of 30 adult females observed, 8 were male-coloured, 12 had tan colour markings, distinctly different from males, and 10 had an intermediate muddy blue-green colouration. Photos: EI Svensson.

Here, we first confirm that female-limited colour polymorphisms in damselflies do typically evolve via the evolutionary origin of male-coloured females, consistent with an advantage of reduced male-mating harassment (20, 32). We then address the two hypotheses outlined above, linking ecological factors to demographic conditions and to the evolution of female-limited polymorphisms in an explicitly causal framework (Fig. 1), through mediation analysis (58). An alternative scenario is that while temperate regions and open landscapes are indeed associated with the occurrence of female-limited polymorphisms, these effects may not be mediated by demographic conditions as we have hypothesized (Fig. 1b). For example, temperature may have a direct effect on male mating effort (59) or male harm upon mating (60), modulating sexual conflict independently of demographic variation. We evaluate the empirical support for these two scenarios and provide evidence in support of a role of high population density in open habitats and temperate regions driving the evolution of male mimicry and female-limited polymorphisms in damselflies. We then follow up this causal inference approach by jointly modelling the evolution of female-limited colour polymorphisms and their environmental predictors, and by reconstructing ancestral population density regimes across the pond damselfly phylogeny. Finally, we investigate the consequences of female-limited colour polymorphisms for the diversification dynamics in this ancient insect group. We show that female polymorphic lineages in damselflies are associated with relatively high diversification rates, similarly to sexually dimorphic lineages and in contrast to sexual monomorphism. These results suggest that sexual interactions brought about by colour signals have important macroevolutionary consequences in damselflies.

## Results

### Macroevolutionary origin of female-limited colour variation

We used a Hidden-State Speciation and Extinction (HiSSE) model to reconstruct the evolutionary history of sex-related colour variation in pond damselflies (Odonata:Coenagrionidae), and to investigate trait-dependent diversification dynamics (see *Diversification consequences of female colour and colour variation*, below). We obtained sex-related colour data for 418 species (out of ∼ 1,400) in the most comprehensive phylogeny of pond damselflies to date (61). Each species was classified as sexually monomorphic (SM), if males and females have the same hues and pattern across the thorax and most abdomen segments, sexually dimorphic (SD), if males and females have distinctly different colour patterns, or female polymorphic (FP), if females occur in two or more distinct colour patterns, one of which is similar to males (see Materials and Methods). The most common character state was sexual dimorphism with 44% of all species, while 28% of taxa were sexually monomorphic and another 28% were female polymorphic.

Our HiSSE analysis did not resolve the colour state of the most recent common ancestor of all pond damselflies, which was inferred as either sexually dimorphic (posterior probability (PP) = 0.565) or as female polymorphic (PP = 0.393; Fig. 2a). Nonetheless, we found moderate support for FP lineages arising with greater probability from SD ancestors than from SM ancestors (difference in transition rate: posterior mean (PM) = 0.004, 95% highest posterior density (HPD) interval = -0.001 -0.010, PMCMC = 0.044; Fig. 2b-c). Stochastic character mapping resulted in a median of 42 gains of female-limited polymorphisms from SD ancestors (95% HPD interval = 4 - 66) versus only three origins of novel female morphs in ancestrally SM lineages (95% HPD interval = 1 - 12). In contrast, female-limited polymorphisms were lost with similar probability to SD and SM descendants (difference in transition rate: PM = 0.001, HPD interval = -0.007 - 0.009, PMCMC = 0.473; Fig. 2b,d).

**Figure 2.**
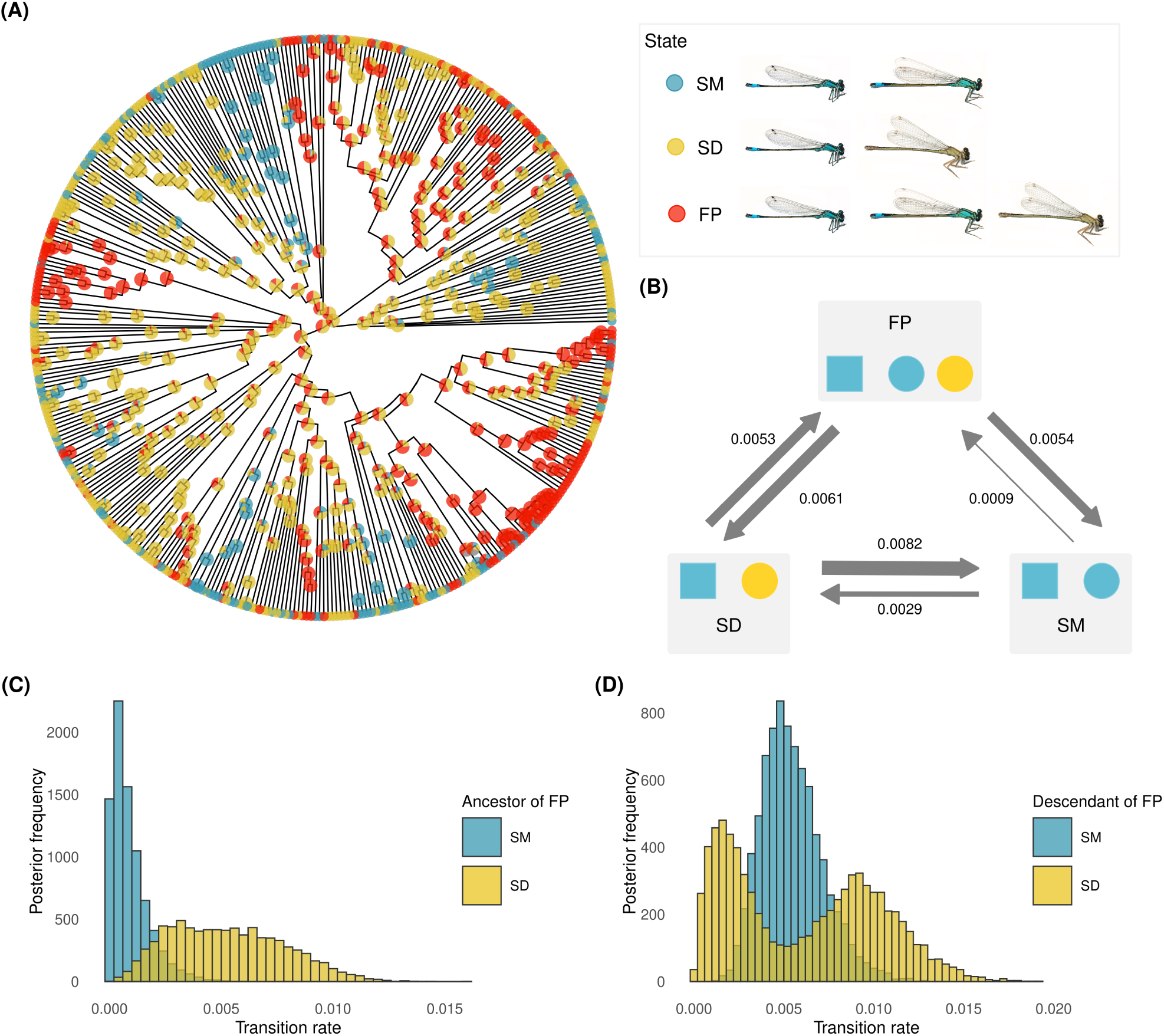
Macroevolutionary history of female-limited colour variation **(A)** Ancestral state reconstruction of sex-related colour states in pond damselflies (family Coenagrionidae). Colours at the tips show colour states in extant taxa. Pies indicate indicate joint conditional posterior probabilities of sex-related colour states in ancestral nodes. **(B)** Summary of transition rates between sex-related colour states in pond damselflies. Arrow width is proportional to estimates of transition rates. Numbers next to arrows represent the mean rate for each transition type. **(C)** Histogram of posterior estimates of the rate of evolution of female-limited polymorphisms, depending on the ancestral colour state. **(D)** Histogram of posterior estimates of the rate of loss of female-limited polymorphisms, depending on the descendant colour state. Abbreviations: FP = female-limited colour polymorphism, SM = sexual monomorphism in colour, SD = sexual dimorphism in colour.

### Ecological factors associated with female-limited colour polymorphisms

Non-ambiguous latitudinal range and habitat openness data were available for 297 of 418 taxa included in our phylogeny (Dataset S1). Using a Bayesian Phylogenetic Mixed Model (BPMM) that adjusts for confounding effects due to shared ancestry of related species (*Model 1*), we found that both latitudinal range and habitat openness predicted the macroevolution of female-limited colour polymorphisms in damselflies. Specifically, female-limited polymorphisms were more common in temperate regions, across both habitat types (temperate vs tropical in open habitats: PM = 0.358, 95% HPD interval = 0.115 - 0.592, PMCMC < 0.001; closed habitats PM = 0.322, 95% HPD interval = 0.060 - 0.653, PMCMC = 0.001; Fig. 3a). FP taxa were weakly associated with more open habitats in tropical regions (PM = 0.056, 95% HPD interval = -0.025 - 0.166, P = 0.077), but not in temperate regions (PM = 0.092, 95% HPD interval = -0.210 - 0.416, P = 0.275; Fig 3a). Unlike female-limited polymorphisms, the occurrence of sexual dimorphism was largely independent from these broad ecological predictors, whereas the presence of sexual monomorphism was associated with tropical ranges and closed habitats (Table S1-S2; Fig. S1).

**Figure 3.**
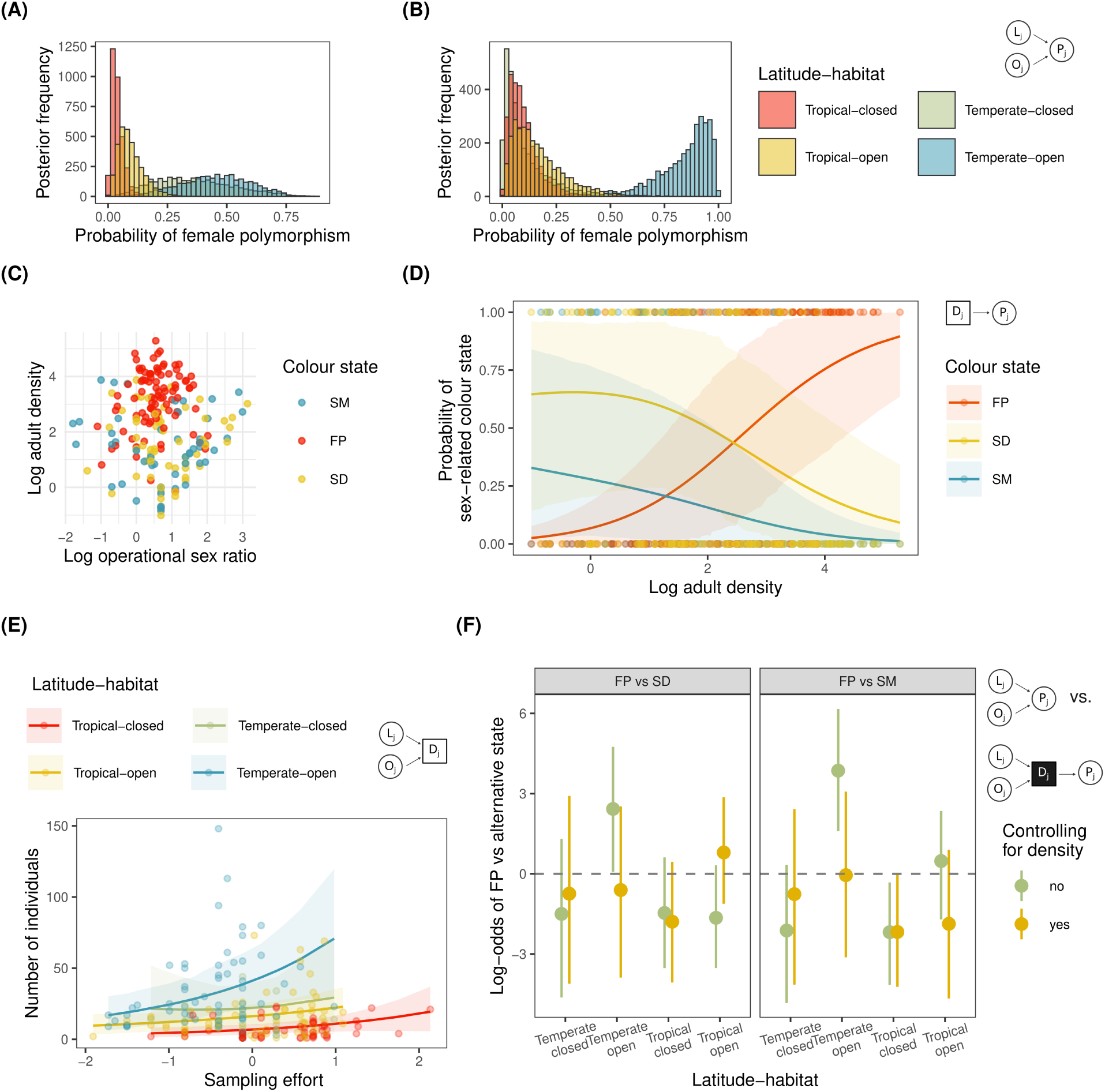
Demography mediates ecological effects on the occurrence of female-limited colour variation. For each panel (except **(C)** which shows raw data), the DAGs show the subset of causal paths from Fig. 1a that were estimated in the corresponding statistical model. The hypothesized mediator (density) is represented by a square and other variables are shown as circles. Analyses in **(A)** and **(B)** are based on the same causal graph. **(A)** Posterior frequency of female-limited polymorphisms depending on latitudinal range and habitat openness and based on the literature-data analysis (*Model 1*). **(B)** Posterior frequency of female-limited polymorphisms depending on latitudinal range and habitat openness and based on the field-data analysis (*Model 2*). **(C)** Operational sex ratio (OSR) and adult population density in the field data set. Each dot represents a population (n = 208) of a total of 80 species. Data are log transformed. **(D)** Predicted probability of each sex-related colour state with increasing (log) population density (*Model 3*). Shaded areas represent 95% HPD intervals. **(E)** Expected number of sampled individuals with increasing sampling effort in each latitude-habitat combination (*Model 5*). Dots show data points and shaded areas represent 95% HPD intervals. Sampling effort is mean centered, so intercept differences represent differences in density between latitude-habitat categories. **(F)** Log-odds ratio of female-limited polymorphisms vs alternative sex-related colour states in models with (*Model 6*) and without (*Model 2*) controlling for (log) adult density. Circles represent posterior means and lines represent 95% HPD intervals. The black-filled density node in the lower DAG indicates that the statistical model adjusted for any predictor effects mediated by adult density. Abbreviations: FP = female-limited polymorphism, SD = sexual dimorphism, SM = sexual monomorphism. Abbreviations in DAGs: L = Latitudinal range, O = habitat openness, D = density,_10_P = female-limited polymorphism.

Female-limited colour polymorphisms occurred with highest probability in open habitats of temperate regions, in both the larger literature-data analysis (*Model 1*, N = 297) and in the analysis including only the taxa surveyed in the field (*Model 2*, N = 80). However, our field-data analysis underestimated the occurrence of female-limited polymorphisms in temperate-closed habitats relative to the literature-data analysis (Fig. 3a-b). Female-limited polymorphisms in the field-data analysis were associated with temperate regions in open-habitat lineages (PM = 0.695, 95% HPD interval = 0.355 - 0.9126, PMCMC = 0.001), but not in closed-habitat lineages (PM = -0.011, 95% HPD interval = -0.215 - 0.305, PMCMC = 0.557). Similarly, female-limited polymorphisms were associated with open habitats in temperate regions in the field-data analysis (open vs closed habitats in temperate regions: PM = 0.466, 95% HPD interval = 0.047 - 0.840, PMCMC = 0.005), but not in the literature-data analysis (Fig. 3a-b). Temperate-closed habitats are the least common habitat type in both data sets, accounting only for ∼5% of the taxa. Thus, limited taxon sampling, especially in the smaller field-collected dataset, likely accounts for the disparity between the two analyses. Finally, within the tropics, we obtained a qualitatively similar result in both analyses, indicating a lower probability of FP in closed habitats. In both the literature- and the field-data analyses the difference between open and closed habitats was of about 5%. However, in the field-data analysis this mean difference was bounded by higher uncertainty (PM = 0.046, 95% HPD interval = -0.151 - 0.327, PMCMC = 0.281).

### Linking ecology, demography, and sexual conflict in damselflies

We used mediation analysis (58) to test whether demographic conditions casually drive the association between female-limited colour polymorphisms and their environmental predictors. This causal inference approach makes three predictions. First, we expect a correlation between the hypothesized demographic mediator and the outcome, in this case the occurrence of female-limited polymorphisms. Second, the demographic mediator should be dependent on the ultimate ecological predictors. Third, there should be an appreciable reduction in magnitude of the parameter estimates of the ultimate ecological drivers once we control for the demographic mediator. In other words, if demographic conditions were held constant, there should be no differences in the occurrence of female-polymorphic taxa across habitats. We focused on two potential mediators of ecological predictors: population density and male-biased OSR at breeding sites (Fig. 3c).

In *Model 3* we found that female-limited colour polymorphisms occurred with increasing probability at higher population densities (slope of log-odds ratio between FP and SM: PM = 1.564, 95% HPD interval = 0.907 - 2.257, PMCMC vs 0 < 0.001; slope of log-odds ratio between FP and SD: PM = 1.094, 95% HPD interval = 0.325 - 1.862, PMCMC vs 0 = 0.004; Fig. 3d). In contrast, a male-biased OSR was not correlated with the incidence of polymorphic females (*Model 4* ; slope of log-odds ratio between FP and SM: PM = 0.008, 95% HPD interval = -0.877 - 0.987, PMCMC vs 0 = 0.994; slope of log-odds ratio between FP and SD: PM = -0.537, 95% HPD interval = -1.529 - 0.362, PMCMC vs 0 = 0.268; Fig. S2-S3). We therefore focused on population density for the subsequent two predictions of a mediator variable.

In *Model 5*, we asked whether the ecological predictors that were associated with female-limited colour polymorphisms (temperate regions and open landscapes) were also associated with the hypothesized mediator: high population density. Once accounting for sampling effort, population densities were indeed higher in temperate-open sites than in all other latitude-habitat categories (Table S3; Fig. 3e, S4). Population densities were lowest in tropical-closed habitats and intermediate in both tropical-open and temperate-closed sites (Table S3; Fig. 3e, S4).

Finally, in *Model 6* we asked if controlling for population density would reduce the expected probability of female-limited colour polymorphism in latitude-habitat categories with higher occurrence of such polymorphisms (temperate-open and, to a lesser extent, tropical-open sites). We found that the log-odds ratio of FP vs. alternative states was markedly reduced in temperate-open sites when we controlled for population density (Fig. 3f), indicating that this demographic variable mediates the macroecological distribution of female-limited colour polymorphisms. In tropical-open sites, accounting for density decreased the log-odds ratio of FP relative to SM, as expected, but increased the log-odds ratio of FP relative to SD (Fig. 3f). This increased log-odds ratio suggests that SD lineages are relatively more common in high-density tropical-open landscapes (Table S1; Fig. S1).

### Macroevolution of ecological and demographic drivers

An assumption of the mediation analysis above is that causality flows from ecological and demographic drivers to the evolution of female-limited colour polymorphism and not the other way around. To inspect this assumption, we used separate discrete-trait models for each ecological factor, and coded both the ecological factor (either latitudinal range or habitat openness) and the female-colour states (female-polymorphic or female-monomorphic) as binary traits. We included a reversible-jump mixture parameter to directly estimate the probability of correlated evolution between the two traits in each model. Evidence for correlated evolution would indicate that evolutionary transitions in at least one trait depend on the character state of the other trait.

Our results show with decisive support (Bayes factor > 100) that the origin of female-limited colour polymorphism depends on the latitudinal range and habitat openness occupied by a lineage (Fig. 4a). As expected, female-limited colour polymorphisms are much more likely to evolve in lineages that already occupy temperate ranges (Fig. 4b; Table S4) and open habitats (Fig. 4c; Table S5). Ancestral state reconstructions under the correlated evolution model between latitudinal ranges and female-colour states provided strong support (PP = 100%) for a most recent common ancestor to all pond damselflies that was female-monomorphic with a tropical range (Fig. S5). Using stochastic character mapping, we estimated 33 (95% HPD interval = 27 - 38) transitions where a shift towards a temperate range predated the evolution of female-limited colour polymorphism, in contrast to a single (95% HPD interval = 0 - 5) transition where a lineage became FP before shifting to a temperate region. Ancestral state reconstructions under the correlated evolution model with habitat openness also inferred a female-monomorphic common ancestor, in this case, occupying open habitats (PP = 100%; Fig. S6). Thus, once a lineage shifts to closed habitats, female-limited colour polymorphisms are less likely to evolve (Fig. 4c). In striking contrast to these findings, latitudinal range and habitat shifts are not strongly dependent on female-colour states (Fig. 4a-c).

**Figure 4.**
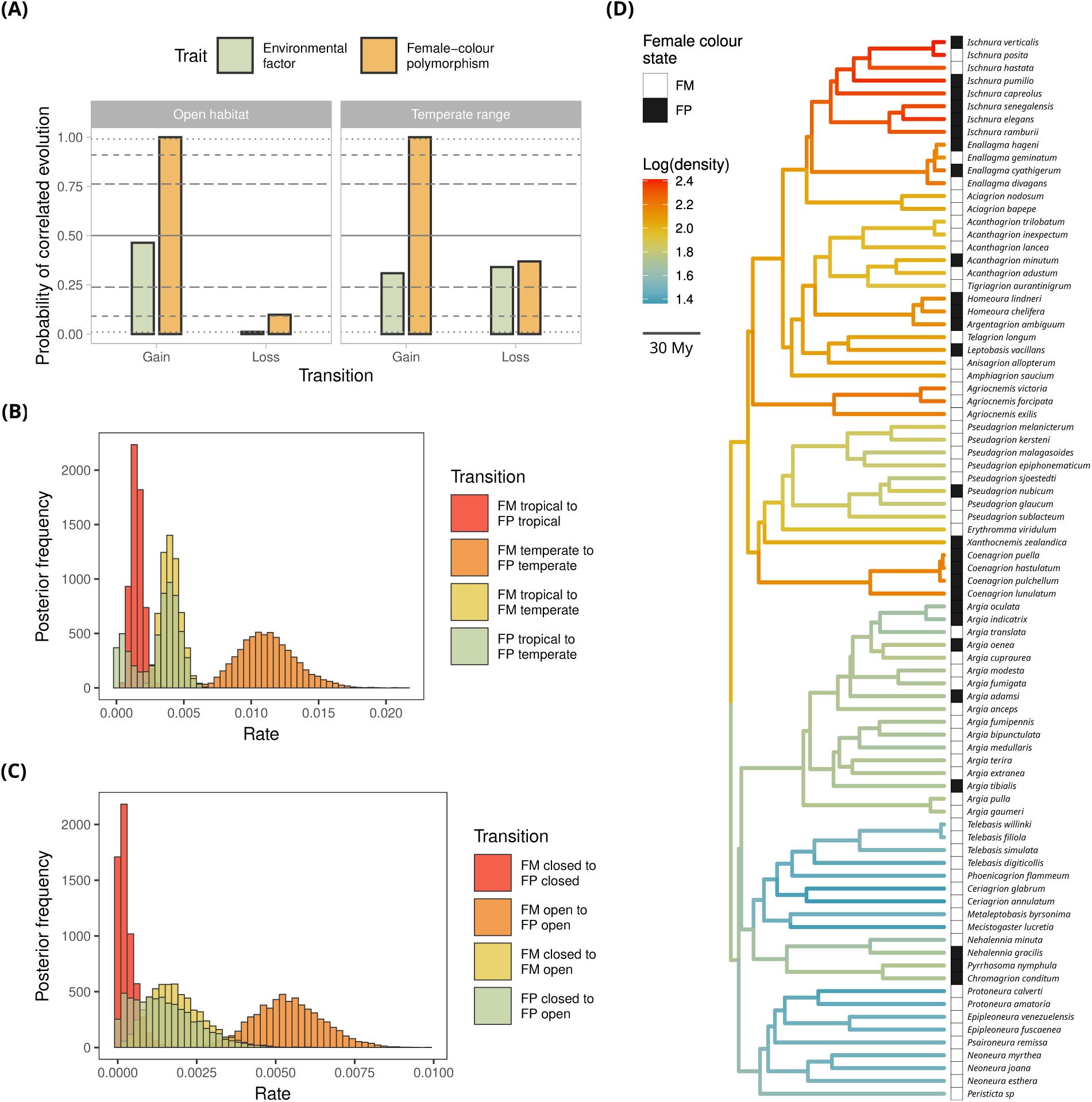
Evolution of ecological and demographic predictors of female-limited colour polymorphisms in pond damselflies **(A)** Probability of correlated evolution between female-limited colour polymorphisms and their two ecological predictors: open habitats and temperate ranges. We used reversible jump Markov Chain Monte Carlo to contrast models in which transition rates are correlated against models with independent transitions. A high probability of correlated evolution indicates strong evidence that an evolutionary transition in one trait depends on the character state of the other. “Gain” transitions refer to the origin of female-limited colour polymorphisms or shifts to open habitats or temperate ranges. “Loss” refers to the fixation of a single female colour morph, or shifts to closed habitats or tropical ranges. The solid line indicates complete uncertainty as to whether traits depend upon each other. Dashed to dotted lines indicate substantial (BF = 3.2), strong (BF = 10), and decisive (BF = 100) evidence in favour of correlated evolution, for high probability values, or in favour of independent evolution, for low probability values. Posterior frequencies of transition rate estimates based on models allowing for correlated evolution between female-limited colour polymorphisms and **(B)** latitudinal range (n = 356 taxa) or **(C)** habitat openness (n = 334 taxa). **(D)** Evolution of adult density optima (*θ*) based on a relaxed Ornstein-Uhlenbeck model. The model was parametrized with a prior expectation of 20 optima shifts across the tree. Results under alternative priors (10 or 40 shifts) are qualitatively similar (Fig. S7-S8). The phylogeny includes only the taxa sampled in the field (n = 83). Multiple density surveys were averaged and density was log-transformed before the analysis.

Because adult density is a continuous variable, measured only for a subset of taxa sampled in field, we are currently restricted in our ability to directly test for correlated evolution between this demographic mediator and female-limited colour polymorphisms, without sacrificing precision and power. Instead, we used a relaxed Ornstein-Uhlenbeck process to model the evolution of adult densities at breeding sites, and inferred branches along the pond-damselfly phylogeny with shifts in density optima. When visualized alongside female-colour states in extant taxa (Fig. 4d, S7-S8), we note that clades with high occurrence of female-limited polymorphisms are associated with positive shifts in density (see Supporting Results in the *SI Appendix*; Table S6).

### Diversification consequences of female colour and colour variation

We estimated the effects of female colour polymorphisms on diversification rates, while controlling for background diversification heterogeneity among clades in a HiSSE model (Fig. S9). Net diversification rates were high and similar in magnitude between female-polymorphic and sexually dimorphic lineages, compared to sexually monomorphic lineages, which had lower diversification rates (Fig. 5; Table S7). However, female-limited colour polymorphisms were inferred as associated with both high speciation and high extinction rates, whereas sexual dimorphism was associated with moderate speciation rates and relatively low extinction rates (Table S7; Fig. S10).

**Figure 5.**
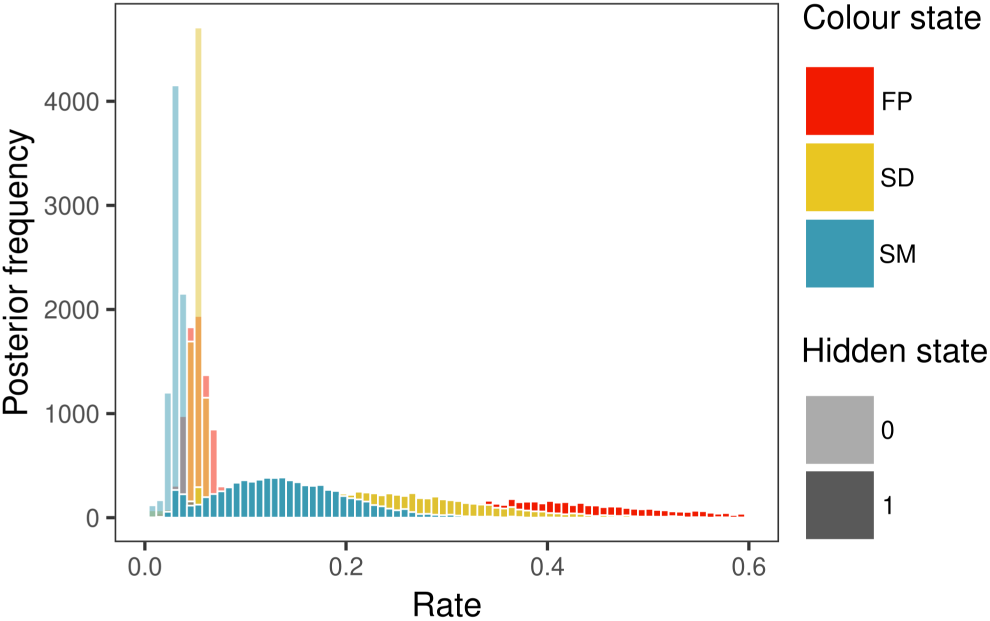
Sex-related colour variation may influence diversification dynamics in pond damselflies. Histograms represent posterior distributions of state-dependent net diversification rates, in a Hidden State-dependent Speciation and Extinction (HiSSE) model. Background heterogeneity in diversification was modeled as a hidden trait with two values (0 = low background diversification, 1 = high background diversification). Abbreviations: FP = female-limited polymorphism, SD = sexual dimorphism, SM = sexual monomorphism.

## Discussion

Our results revealed a demographic link between ecology and sexual conflict in the evolution of female-limited colour polymorphisms in pond damselflies (Fig. 3). Temperate and open landscapes across the globe have the highest densities of reproductive individuals, which in turn results in a higher incidence of female-polymorphic taxa. In the tropics, a similar pattern is observed in open landscapes, where damselfly species typically occur at higher densities than in the forest, and female-limited colour polymorphisms are also more common than in closed habitats. These results are in line with theoretical expectations and intraspecific studies in several different insect taxa, that have argued or showed that increased population density elevates the intensity of sexual conflict over mating, by accelerating the rate of pre-mating interactions (20, 38, 45, 46, 62). Moreover, our findings suggest that female-limited polymorphisms have arisen predictably and repeatedly across the evolutionary history of pond damselflies (Fig. 2, S5-S6). The multiple independent evolutionary origins of female-limited polymorphisms are associated with the same social selection regime (sexual conflict over mating) shown to maintain these polymorphisms within sympatric populations (29–33).

Bayesian phylogenetic mixed models (Fig. 3a-b) and correlated evolution analyses (Fig. 4a-c) jointly evidenced that temperate ranges and open habitats promote the evolution of female-limited colour polymorphisms. We also inferred evolutionary shifts towards higher adult densities in clades where the majority of species are female polymorphic. These findings are causally integrated through our mediation analysis (58); if there were no changes to adult density in response to latitudinal and habitat shifts, the relationship between ecological factors and female-limited colour polymorphisms would break down (Fig. 3f). Mediation analysis makes three assumptions that warrant consideration. First, we assumed there are no missing confounders in the relationship between adult density and female-limited colour polymorphisms. If there was an unmeasured variable independently driving both our hypothesized mediator and outcome, estimation could be biased. Predation pressure might directly reduce density (63) and also reduce male-mating harassment, if individuals become less active to avoid risk (64). Thus, a model that incorporates the effects of predation could challenge these interpretations. Second, there should be no interaction, such that density mediates the occurrence of female-limited colour polymorphisms in only some latitude-habitat combinations. A compelling test of this assumption would require additional experimental or field data on the effects of density under varying ecological conditions. Third, we assumed that the direction of causality is not mistaken: the preexistence of female-limited colour polymorphisms should not facilitate habitat or range shifts. Our correlated evolution analyses strongly support the validity of this assumption (Fig. 4a-c).

In the present study, male-biased operational sex ratio (OSR) was not associated with any sex-related colour state (Fig. S2-S3). Foundational theory by Emlen and Oring (65) predicted that a male-biased OSR should increase the opportunity for sexual selection on males, a pattern supported by large-scale comparative evidence (40). We thus expected male-biased populations to be associated with a higher probability of sexual dimorphism. While a male-biased OSR may increase the opportunity for sexual selection on damselfly males as in other taxa, this selection regime may not translate into greater sexual dimorphism in colour. In territorial damselflies, OSR is often male-biased, with males defending small patches along breeding sites and females visiting these sites only when reproductively available (e.g. 66). In these species, females often select mates based on the quality of oviposition sites in their territories and male-male competition for suitable sites might result in sexual dimorphism in size, instead of colour (e.g. 67). Alternatively, relevant effects of OSR on sex-related colour variation may have been obscured by measurement error arising from having sampled one or a few snapshots of natural populations. OSR can fluctuate rapidly within mating seasons (68), and vary across microhabitats (69). Finally, even if OSR is estimated without error, it remains an incomplete measure of the opportunity for sexual selection, especially when the strength of competition for mates also depends on other factors (70). For instance, in the extreme case where density is so high that males are unable to effectively monopolize mates, selection for investment in competitive traits might be weakened even is the OSR is male-biased (70).

The absence of a clear effect of OSR on female-limited colour polymorphisms in the present study was also at odds with expectations from previous experimental studies, in which male-biased OSR has been found to increase the intensity of sexual conflict over mating (49, 50). We therefore expected that female-limited colour polymorphisms would increase in frequency with more male-based OSR, as well as with higher adult density. Unlike experimental and modelling studies in which the effects of these demographic factors can be neatly teased apart, natural populations may only occupy a viable subset of all possible parameter combinations. Even if the combination of high adult density and male-biased OSR would likely exacerbate the rate of male-mating harassment, such a demographic environment may be too hostile to female fitness be sustained in the long term (71). In our field data set, only 2.4% of all populations (five populations of four species) were in the top 20% of both adult density and male-biased OSR (Fig. S2). While all of these populations were polymorphic, as expected, our data suggest that even with the evolution of female-limited polymorphisms, combinations of demographic conditions that are extremely adverse for females may be rare, as populations inhabiting such environments might suffer from elevated extinction risk (72, 73).

We hypothesized that environmental conditions in the temperate zone and open habitats promote the synchronous emergence of adults and their permanence at breeding sites, in turn leading to high adult densities and male-male scramble competition that could override female choice (Fig. 1a). When each female is likely to encounter multiple males, males benefit more from repeated pre-mating attempts and females suffer a greater harassment cost of being single (62). As a result, female choosiness is reduced and overall mating rates increase. Under these circumstances of high population density and female costs from both excessive mating and pre-mating harassment, the emergence of a novel female morph that avoids excessive male attention could be favoured by natural selection. Such a morph could then increase in frequency and subsequently be maintained at some intermediate level by negative frequency-dependent selection (29–31). In line with this idea, our HiSSE model revealed that sexual dimorphism was the most likely ancestral state to female-limited colour polymorphisms in pond damselflies (Fig. 2). These results suggest that female-limited colour polymorphisms typically evolve through the origin of a novel male-coloured female morph in a sexually dimorphic background, consistent with this female morph being a male mimic.

We note that non-selective genetic mechanisms might operate alongside a selective advantage of male mimicry at high density. Female-limited colour morphs in several orders of insects differ from each other in the expression of regulatory elements that interact with sex-determination loci (74–76). Thus, while the origin of female morph with a novel colour pattern likely requires evolution of sex-limited expression, a novel male-coloured female morph can arise via any mutation that disrupts pre-existing sex-specific regulation. Male-coloured females could therefore arise at higher rates than other novel morphs, even without any fitness advantage. Yet, such neutral female-limited polymorphisms are likely short-lived and eventually eroded by genetic drift unless maintained by some form selection (77). Consistent with such a scenario of mutation bias, the HiSSE model also revealed a higher rate of transition from sexual dimorphism to sexual monomorphism than *vice versa* (Fig. S11).

Female polymorphic lineages have higher diversification rates compared to sexually monomorphic lineages, but not relative to sexually dimorphic lineages (Fig. 5). Sexual selection, for which sexual dichromatism is often used as a proxy, has been found to promote speciation in some clades (78, 79) but not in others (80, 81). There is evidence that sexual conflict can also promote diversification (82), although this outcome might depend on demographic conditions, with large populations more likely fostering sympatric speciation (83). Rapid diversification in female-polymorphic lineages is also consistent with “morphic speciation”, whereby different morphs are lost due to genetic drift in allopatric populations, promoting reproductive isolation (84, 85).

We caution that birth-death models, have increasingly recognized identifiability limitations. A potentially infinite number of parameter combinations are equally likely to generate the diversity pattern captured by a molecular phylogeny (86). Hidden states in HiSSE models have a similar effect; their parametrization generates congruent classes of trait-dependent and trait-independent diversification scenarios that could have given rise to the same observed data (87). Therefore, diversification results in this study should be considered tentative. With these reservations in mind, the HiSSE analysis suggested that female-polymorphic lineages diversify rapidly despite a trend towards increased extinction (Fig. S10). Even if correct, elevated extinction rates may not be a consequence of the polymorphism per se, but could be driven by the same process of sexual conflict that promotes the evolution of polymorphisms in the first place. Sexual conflict over mating rates can increase population extinction risk through its negative effects on female fitness, an idea supported both in theory (71), and by some empirical evidence (88, 89).

### Conclusions

Ecologists have long acknowledged the power of density-dependent natural selection, promoting genetic and phenotypic divergence (90, 91). While the role of ecological factors on the intensity and outcome of sexual conflict has recently attracted research interest (92, 93), the mechanisms for such ecological effects have rarely been examined (e.g. 60). Here, we have shown that population density causally links environmental heterogeneity to the phylogenetic distribution of female colour polymorphisms in damselflies (Fig. 3). High population density and elevated sexual conflict at high latitudes and in open habitats has repeatedly favoured the evolution of female-limited colour polymorphisms, in support of the hypothesis that male mimicry is an evolved female response to male mating harassment (Fig. 1a). Taken together with previous microevolutionary studies in damselflies (20, 29–33), these results indicate that sexual conflict plays an important role both in the evolutionary origin and in the maintenance of female-limited colour polymorphisms. Our results reveal how phylogenetic comparative analyses can be enriched by incorporating predictions from microevolutionary studies to reveal the ecological drivers and macroevolutionary consequences of sexual conflict.

## Methods

### Field work and collection of comparative phenotypic data

We obtained comparative phenotypic data from field surveys, literature, and online database searches and examination of museum specimens. Field surveys are described in detail below. For literature data, we mined field guides, species descriptions, and primary scientific literature, including taxonomic revisions and regional checklists. We also obtained data from online resources managed or contributed by expert odonatologists, and from descriptions accompanying museum specimens, housed in the Florida State Collection of Arthropods (https://iodonata.updog.co/index_files/fscaodon.html) and in the New Zealand Arthropod Collection (https://www.landcareresearch.co.nz/resources/collections/nzac). All comparative data and references to the sources used for each species are available as Supporting Information (Dataset S1-S2).

For each species, we recorded the following variables whenever available: sex-related colour variation (see below), latitudinal range (tropical or temperate), habitat openness, population density and operational sex ratio. Taxa (n = 418) were classified as sexually monomorphic (SM), sexually dimorphic (SD) or female polymorphic (FP), depending on whether females are male-coloured, markedly different from males, or if multiple discrete female colour morphs have been documented. Females were classified as male-coloured if they exhibit similar hues (e.g. black, blue, green, yellow, red, brown) and similar patterning (i.e. plain or patterned) as males in both their thorax and most of the length of their abdomen (segments 1-7). Females were otherwise classified as different from males in colour. See the *SI Appendix* for details on female-colour classification.

Latitudinal distribution data were taken from Willink et al. (61), where species distributions were recorded as presence or absence from administrative areas, namely countries and states, provinces, or territories. The resulting distribution ranges were then classified as warm-adapted (tropical) or temperate or cold-adapted (temperate) based on the biome classification of the areas they occupy. Species that covered both latitudinal ranges were treated as missing data (n = 83). Breeding habitats were categorized as open or closed based on the vegetation type and cover. In the field, we classified habitats as open if the prevailing vegetation were grasses and shrubs, and as closed for species collected within the forest understory. Similarly, when gathering habitat openness data from the literature we considered descriptions including the words “open”, “marsh”, “bog”, and “grassland” or “grasses” as indicating open habitats, and descriptions including the words “closed”, “shaded”, “wooded”, and “forest” as indicating closed habitats. Here too, species with mixed habitat descriptions in the literature (n = 106) were treated as ambiguous, unless the literature source explicitly referred to one habitat type being more common than the other.

Density and operational sex ratio data were collected only for species surveyed in the field, and for one additional species (*Xathocnemis zealandica*, in New Zealand) that was surveyed in a previous study using comparable field methods as ours (94). We conducted field surveys in four continents and six countries (Argentina, Costa Rica, Guyana, Cameroon, Sweden, Cyprus, and USA) in order to sample a wide ecological and phylogenetic range of taxa. In each country, we surveyed 9-29 breeding sites, for an average of 2.6 netting hours per site (sd = 3.2 h, min = 12 min, max = 16.3 h). At each site, one to three researchers captured damselflies as they were encountered along a transect, during a time-recorded capture session. All captured specimens were subsequently identified to species, sex, and colour morph when applicable. If more than 8 individuals per species were collected, these timed surveys were used to calculate an index of population density as the number of individuals caught per researcher per hour, and an index of operational sex ratio (OSR), defined as the ratio of males per females at the breeding sites. While the population density index should reflect the overall intensity of intra-specific interactions, the OSR is often taken as a measure for the opportunity of sexual selection (40). In some species, only members of one sex were observed (typically males). In order to include these sex-biased populations in our OSR analysis, we estimated OSR in all taxa as (number of males + 1)/(number of females + 1), providing a lower boundary estimate of the extent of male bias. Restricting the analysis to only populations in which both males and females were observed did not qualitatively change our results.

### Estimating transition rates and diversification dynamics

We addressed two questions about the evolutionary history of female-limited colour variation. First, we asked which ancestral state, sexual monomorphism (SM) or sexual dimorphism (SD), more often precedes the evolution of female-limited colour polymorphism (FP). A higher transition rate from SD to FP would indicate that that female-limited polymorphisms typically evolve from sexually dimorphic ancestors, and through the origin of a novel male-like female phenotype, consistent with the male-mimicry hypothesis (95). Next, we asked whether there was any bias in the female morph that persists if and when female-limited colour polymorphisms are lost. To answer these questions we inferred the transition rates between sex-related colour states (SM, SD, and FP) across the phylogeny of pond damselflies, using a Hidden-State Speciation and Extinction (HiSSE) model. We chose to model the evolution of sex-related colour variation while accounting for heterogeneity in diversification rates because ignoring such diversification asymmetries can result in an overestimation of transitions towards diversification-enhancing states (96), and our preliminary analysis suggested that diversification dynamics differed between character states (see Results).

We implemented the HiSSE analysis in RevBayes v. 1.1.1 (97). This model allows for character-dependent and background heterogeneity in diversification rates, by modelling the diversification effects of an unobserved trait. For convenience, we assumed that the hidden character influencing background heterogeneity in diversification had two alternative states. We based the analysis on the maximum *a posteriori* (MAP) tree from Willink et al. (61), and explicitly considered the proportion of missing taxa as randomly distributed across the tree. Information on priors and MCMC simulations for HiSEE and all subsequent analyses are available in the *SI Appendix*.

### Ecological effects and demographic mediation of sex-related colour variation

We investigated if the evolution of female-limited colour polymorphisms could be predicted by environmental factors, and if these effects are mediated by demographic conditions (see Introduction). In *Model 1*, we explored the relationship between the presence of female-limited colour polymorphisms and two ecological factors: latitudinal range, a proxy for the duration and intensity of the mating season, and habitat openness, as a proxy for the isolation of reproductive habitats and the potential for females to escape male mating harassment through local movements. We conducted this and subsequent analyses in R v 4.2.2 (98), using Bayesian phylogenetic mixed-effect models (BPMM) implemented with the package *MCMCglmm* v. 2.33 (99). In all BPMM models, we adjusted confounding effects due to shared evolutionary ancestry, by fitting the phylogenetic variance–covariance matrix, derived from the phylogenetic tree, as a random effect. Sex-related colour state (the response variable) was coded as a multinomial variable with the same three categories as above: sexual monomorphism (SM), sexual dimorphism (SD), and female-limited colour polymorphism (FP). The fixed latitude and habitat predictors had two levels each: ‘tropical’ and ‘temperate’, and ‘open’ and ‘closed’, respectively, and with ambiguous taxa coded as missing data and thus excluded from the analyses.

In our subsequent BPMMs (*Model 2* - *Model 6*), we used only the data from species which we sampled ourselves in the field, or for which comparable demographic data was available from published studies (Dataset S2). In *Model 2*,* we specified the same model as outlined above and corroborated that the qualitative effects of latitudinal range and habitat openness were similar with the reduced data set. Then, we explicitly tested if the effects of these ecological factors on the evolution of sex-related colour variation are mediated by demographic conditions, specifically, local density and sex ratio at breeding sites. To do this, we first quantified the effect of the demographic variables on the probability of female-limited colour polymorphisms, using multinomial BPMMs and observed densities (log number of individuals per hour, *Model 3*) and operational sex ratios (log number of males per female, *Model 4*) as predictors. Because in many cases different species were surveyed in the same breeding sites and some species were surveyed in multiple breeding sites, we included the breeding site identity as an additional random effect in all BPMMs using field data.

After observing that only one of the demographic variables, population density, predicts the occurrence of female-limited colour polymorphisms (see Results), we conducted two additional tests to determine whether population density is a mediator of broad-scale ecological factors (latitudinal range and habitat openness). In *Model 5*, we tested for differences in population density between habitats and latitudinal ranges, modelling the number of individuals caught per sampling event with a Poisson distribution, and using the sampling effort (catching time) as a mean-centered covariate.

As the final prediction in mediation analysis (58), we asked if ecological predictors become less informative when population density is held constant. If open and temperate habitats indeed have more female-polymorphic lineages because they promote higher adult densities at breeding sites, we would then expect that once we know the adult density of a population, knowing its habitat type and latitudinal range should not change our expectation of finding female-limited morphs. To test this hypothesis in *Model 6*, we included both ecological factors and population density as predictors in a single multinomial model. We report the log-odds ratio of FP vs. each of the two alternative states (SM and SD) for each latitude-habitat combination, with and without accounting for population density. We predicted that the log-odds ratio in latitude-habitat categories that are associated with female-limited polymorphisms should drop to the null expectation (the log ratio of relative frequencies) when density is held at a low value (log density = 0, or 1 adult per hour).

### Correlated evolution of ecological factors and female-colour states

We used correlated evolution models in RevBayes v. 1.1.1 (97), to determine if evolutionary transitions in female-colour states depend upon latitudinal range and habitat openness, but not vice versa. To test this hypothesis, we included a reversible-jump mixture that visits correlated evolution and independent evolution model parametrizations. Thus, this parameter indicated the probability of favouring a correlated evolution model, and was used to compute the raw Bayes Factor for the strength of evidence in favour of a correlated evolution scenario. We conducted two separate analysis for the latitudinal range and habitat openness predictors of female-limited colour polymorphisms. This modelling framework requires binary traits, so for these analyses we re-coded sexually monomorphic and sexually dimorphic lineages as female-monomorphic (FM). We based these analyses on the same literature-based data as *Model 1* in our BPMM approach. Because we investigated the two ecological predictors in separate analyses, here we assessed data completeness and ambiguity for only the modelled ecological factor, resulting in 356 present-day taxa for the correlated evolution analysis including latitudinal range, and 334 taxa for the correlated evolution analysis including habitat openness.

### Estimating evolutionary shifts in adult density optima

We used a relaxed Ornstein-Uhlenbeck (OU) process to model the evolution of population densities across the pond damselfly phylogeny and infer ancestral shifts in population density optima. As input data, we used the log-transformed adult densities surveyed in natural populations of 83 species of pond damselflies. If multiple populations were sampled for the same species, we used the average density across populations as our density estimate. The relaxed OU model includes the three parameters of a standard OU model: the stochastic evolution rate (*σ*^2^), the pull to the optimum (*α*), and the optimum value at the root of the tree (*θ*). Additionally, the relaxed OU model includes a reversible-jump mixture distribution governing the probability of shifts in population density optima on each branch of the tree. This distribution was parametrized with the prior expectation on the total number of shifts across the tree. Because we had no knowledge of how many times adult densities could have shifted, we ran separate models with expectations of 10, 20, and 40 shifts in total.

## Acknowledgements

We are grateful to all the field assistants who helped with the damselfly population surveys, especially Robin Pranter, Hanna Bensch, Yassin Tschinda, Geovanna Rojas, and Emmanuel Venegas. These surveys were conducted with research permits from the Guyana Environmental Protection Agency (Reference No. 010715 BR001), the Administración de Parques Nacionales, Argentina (Ref No. NEA 401), from the Cameroon Ministry of Scientific Research and Innovation and Ministry of Forestry and Wildlife (Ref. No. 0000034), and from the Sistema Nacional de Áreas de Conservación, Costa Rica (M-P-SINAC-ACAT-083-2019, SINAC-ACC-PI-R-122-2019, R-SINAC-PNI-ACLAC-065-2019, SINAC-ACOSA-DT-PI-R-004-2020). We acknowledge the logistic support from the Iwokrama International Centre for Rain Forest Conservation and Development, Guyana, the Karanambu Trust, Guyana, the National Park Administration, Argentina, and the Congo Basin Institute, Cameroon. The Swedish National Infrastructure for Computing (SNIC) provided computational resources for comparative analyses and the Centre for Scientific and Technical Computing at Lund University (LUNARC) provided technical assistance. Funding for this study was provided by research grants from The Swedish Research Council (VR: grants no. 2016-03356 and 2020–03123), the Swedish Foundation for International Cooperation in Research and Higher Education (STINT) and Erik Philip Sörensens Stiftelse to E.I.S. B.W. was supported by a “Faculty for the Future” scholarship from the Schlumberger Foundation and a international Postdoc Grant from the Swedish Research Council (VR: grant no. 2019-06444). We thank two anonymous reviewers for the valuable suggestions and feedback.

## Author contributions

**EIS** and **BW** planned the study and collected the field data. **TATH** and **BW** compiled the literature data. **BW** designed and conducted the phylogenetic comparative analyses. **BW** wrote the manuscript with input from **EIS** and **TATH**.

