## Supporting Information Appendix for "Ecology and sexual conflict drive the macroevolutionary dynamics of female-limited colour polymorphisms"

B. Willink[1*†] T. A. T. Ho[3] E. I. Svensson[4]

2025-06-12

[1] Department of Entomology, Cornell University, Ithaca, NY 14853, USA

[2] Department of Biology, Aarhus University, Aarhus C 8000, Denmark

[3] Department of Biology, Division for Biodiversity and Evolution, Ecology Building, Lund University, Lund 223-62, Sweden

**Keywords**: directed acyclical graphs, diversification, male-mimicry, phylogenetic comparative methods, population density

**References** **21**

### Comparative Data

#### Preview of Dataset S1: Literature data

**Dataset S1**. Latitudinal range, habitat, and colour data for species of pond damselflies (family Coenagrionidae), included in our phylogeny. Here, we show the first ten taxa in alphabetical order. The complete data set is available as a supporting tab-separated file “SupportingTableS1.tsv”. “Latitude” is a discrete variable with two states, for Tropical (Trp) and Temperate (Tmp) ranges, respectively. “Habitat” refers to whether damselflies breed in open (O) landscapes, such as meadows, bogs, and grasslands, or in closed environments (C), such as forests. Latitudinal ranges spanning both hemispheres and broad habitat preferences are treated as missing data. For each species, we then report the occurrence of females that are markedly different from males (heteromorphic females; “HFemale”), and females with similar colour patterns as males (andromorphic females; “AFemale”). If both types of females are known to occur, the species is coded as an occurrence of a female “Polymorphism”. For our statistical analysis, we collated female colour information into a single variable (“FemState”), with states 0 = sexual monomorphism (andromorphic females only), 1 = female polymorphism (andromorphic and heteromorphic females co-occur), and 2 = sexual dimorphism (heteromorphic females only).

To determine the occurrence of alternative female types we scored the colour pattern of males and females from available images and specimens. The dorsal colour pattern of males (Male), andromorphic females (“AFem”) and heteromorphic females (“HFem”) was scored separately for the thorax (“T”), abdomen segments 1-7 (“S17”), and abdomen segments 8-9 (“S89”). For each segment, we scored the presence of a pattern (“P”). The presence of a pattern implies that two colours (“C1” and “C2”) for a given body region are different, whereas a plain-coloured body region has the same colour value for the two colour columns. This results in nine columns with composite names, for each sex and female type. For example, the column “AFemS17C1” refers to the colour value of the first colour in abdomen segments 1-7 for adromorphic females. The colour values are BLA = black, BLU = blue, BRO = brown, GRE = green, GRY = grey, IBLA = iridescent black, IGE = iridescent green, IR = iridescent red, O = orange, R = red, V = Violet, W = white, Y = yellow.

Finally, for each species, we list the resources from ecological and colour data were obtained. These include books (primarily field guides), research articles (usually taxonomic studies or reports from surveys), IUCN red list assessments, other reputable online resources (such as iNaturalist), and whether we observed this species in the field and museum collections. Full references are available through the supporting file “Table_S1.bib”. Collection names are: FSCA = Florida State Collection of Arthropods, SI = Smithsonian Institute, NZAC = New Zealand Arthropod Collection.

| Taxon | Latitude | Habitat | HFemale | AFemale | Polymorphism | FemState | ... |
|---|---|---|---|---|---|---|---|
| *Acanthagrion_adustum* | Trp | O | 0 | 1 | 0 | 0 | ... |
| *Acanthagrion_amazonicum* | | | | | | | ... |
| *Acanthagrion_apicale* | | | | | | | ... |
| *Acanthagrion_cuyabae* | Trp | O | 0 | 1 | 0 | 0 | ... |
| *Acanthagrion_floridense* | Trp | O | 0 | 1 | 0 | 0 | ... |
| *Acanthagrion_fluviatile* | | | | | | | ... |
| *Acanthagrion_gracile* | | | | | | | ... |
| *Acanthagrion_inexpectum* | Trp | O | 0 | 1 | 0 | 0 | ... |
| *Acanthagrion_lancea* | Trp | O | 0 | 1 | 0 | 0 | ... |
| *Acanthagrion_minutum* | Trp | O | 1 | 1 | 1 | 1 | ... |
| ... | ... | ... | ... | ... | ... | ... | ... |

| Taxon | MaleTP | MaleTC1 | MaleTC2 | MaleS17P | MaleS17C1 | MaleS17C2 | MaleS89P | MaleS89C1 | MaleS89C2 | ... |
|---|---|---|---|---|---|---|---|---|---|---|
| *Acanthagrion_adustum* | 1 | BLA | O | 0 | BLA | BLA | 1 | BLA | O | ... |
| *Acanthagrion_amazonicum* | | | | | | | | | | ... |
| *Acanthagrion_apicale* | 1 | BLA | O | 0 | BLA | BLA | 1 | BLA | BLU | ... |
| *Acanthagrion_cuyabae* | 1 | BLA | BLU | 0 | BLA | BLA | 1 | BLA | BLU | ... |
| *Acanthagrion_floridense* | 1 | BLA | BLU | 0 | BLA | BLA | 1 | BLA | BLU | ... |
| *Acanthagrion_fluviatile* | | | | | | | | | | ... |
| *Acanthagrion_gracile* | 1 | BLA | BLU | 0 | BLA | BLA | 1 | BLA | BLU | ... |
| *Acanthagrion_inexpectum* | 1 | BLA | BLU | 0 | BLA | BLA | 1 | BLA | BLU | ... |
| *Acanthagrion_lancea* | 1 | BLA | BLU | 0 | BLA | BLA | 1 | BLA | BLU | ... |
| *Acanthagrion_minutum* | 1 | BLA | BLU | 1 | BLA | BLU | 1 | BLA | BLU | ... |
| ... | ... | ... | ... | ... | ... | ... | ... | ... | ... | ... |

| Taxon | AFemTP | AFemTC1 | AFemTC2 | AFemS17P | AFemS17C1 | AFemS17C2 | AFemS89P | AFemS89C1 | AFemS89C2 | ... |
|---|---|---|---|---|---|---|---|---|---|---|
| *Acanthagrion_adustum* | 1 | BLA | O | 0 | BLA | BLA | 0 | BLA | BLA | ... |
| *Acanthagrion_amazonicum* | | | | | | | | | | ... |
| *Acanthagrion_apicale* | | | | | | | | | | ... |
| *Acanthagrion_cuyabae* | 1 | BLA | BLU | 0 | BLA | BLA | 0 | BLA | BLA | ... |
| *Acanthagrion_floridense* | 1 | BLA | BLU | 0 | BLA | BLA | 1 | BLA | BLU | ... |
| *Acanthagrion_fluviatile* | 1 | BRO | BLU | 0 | BLA | BLA | 1 | BLA | BLU | ... |
| *Acanthagrion_gracile* | | | | | | | | | | ... |
| *Acanthagrion_inexpectum* | 1 | BLA | BLU | 0 | BLA | BLA | 1 | BLA | BLU | ... |
| *Acanthagrion_lancea* | 1 | BLA | BLU | 0 | BLA | BLA | 1 | BLA | BLU | ... |
| *Acanthagrion_minutum* | 1 | BLA | BLU | 1 | BLA | BLU | 1 | BLA | BLU | ... |
| ... | ... | ... | ... | ... | ... | ... | ... | ... | ... | ... |

| Taxon | HFemTP | HFemTC1 | HFemTC2 | HFemS17P | HFemS17C1 | HFemS17C2 | HFemS89P | HFemS89C1 | HFemS89C2 | ... |
|---|---|---|---|---|---|---|---|---|---|---|
| *Acanthagrion_adustum* | | | | | | | | | | ... |
| *Acanthagrion_amazonicum* | | | | | | | | | | ... |
| *Acanthagrion_apicale* | | | | | | | | | | ... |
| *Acanthagrion_cuyabae* | | | | | | | | | | ... |
| *Acanthagrion_floridense* | | | | | | | | | | ... |
| *Acanthagrion_fluviatile* | | | | | | | | | | ... |
| *Acanthagrion_gracile* | | | | | | | | | | ... |
| *Acanthagrion_inexpectum* | | | | | | | | | | ... |
| *Acanthagrion_lancea* | | | | | | | | | | ... |
| *Acanthagrion_minutum* | 1 | BLA | BRO | 1 | BLA | BRO | 1 | BLA | BRO | ... |
| **...** | **...** | **...** | **...** | **...** | **...** | **...** | **...** | **...** | **...** | **...** |

| Taxon | Book | Article | IUCN | Online | Field | Collection |
|---|---|---|---|---|---|---|
| *Acanthagrion_adustum* | Heckman (2008) | | | | Guyana | FSCA |
| *Acanthagrion_amazonicum* | Heckman (2008) | | | | | FSCA |
| *Acanthagrion_apicale* | Heckman (2008) | | | | | FSCA |
| *Acanthagrion_cuyabae* | Heckman (2008) | | von Ellenrieder (2009a) | https://inaturalist.ca/taxa/ 92996-Acanthagrion-cuyabae/ | | |
| *Acanthagrion_floridense* | Heckman (2008) | | Lozano and Muzon (2020a) | https://inaturalist.ca/taxa/ 197549-Acanthagrion-floridense/ | | FSCA |
| *Acanthagrion_fluviatile* | Heckman (2008) | von Ellenrieder et al (2017) | | | | FSCA |
| *Acanthagrion_gracile* | Heckman (2008) | | | https://inaturalist.ca/taxa/ 197547-Acanthagrion-gracile/ | | |
| *Acanthagrion_inexpectum* | Heckman (2008), Paulson and Haber (2021) | | von Ellenrieder (2009b) | https://inaturalist.ca/taxa/ 92998-Acanthagrion-inexpectum/ | | FSCA |
| *Acanthagrion_lancea* | Heckman (2008) | | | https://inaturalist.ca/taxa/ 197553-Acanthagrion-lancea/ | | |
| *Acanthagrion_minutum* | Heckman (2008) | von Ellenrieder et al (2017) | | | Guyana, Argentina | FSCA |
| **...** | **...** | **...** | **...** | **...** | **...** | **...** |

#### Preview of Dataset S2: Field data

**Dataset S2**. Ecological, demographic, and colour data for species of pond damselflies (family Coenagrionidae) sampled in the field. Here, we show the first twenty taxa in alphabetical order. The complete data set is available as a supporting tab-separated file “SupportingTableS2.tsv”. We report the “Country” and “Site” for each observation of each species. “Lat” is a discrete variable with two states, for Tropical (Trp) and Temperate (Tmp) ranges, respectively. “Habitat” refers to whether damselflies breed in open (O) landscapes, such as meadows, bogs, and grasslands, or in closed environments (C), such as forests. “FemState” indicates the sex-related colour state as in Table S1, with values: 0 = sexual monomorphism (andromorphic females only), 1 = female polymorphism (andromorphic and heteromorphic females co-occur), and 2 = sexual dimorphism (heteromorphic females only). During each field visit, we recorded the total number of females (“Female”) and males (“Male”) and the total catching effort in minutes (“Time”). These three variables were used to estimated the operational sex ratio (OSR) and adult population density as explained in the Methods.

| Taxon | Country | Site | Lat | Hab | FemState | Female | Male | Time |
|---|---|---|---|---|---|---|---|---|
| *Acanthagrion_adustum* | Guyana | Iwokrama Esequibo River | Trp | O | 0 | 4 | 15 | 200 |
| *Acanthagrion_apicale* | Guyana | Iwokrama road stream | Trp | C | | 0 | 2 | 140 |
| *Acanthagrion_apicale* | Guyana | Iwokrama road ponds | Trp | C | | 0 | 5 | 182 |
| *Acanthagrion_inexpectum* | Guyana | Iwokrama road ponds | Trp | O | 0 | 0 | 6 | 145 |
| *Acanthagrion_inexpectum* | Costa Rica | Pond at La Gamba Rainforest Lodge | Trp | O | 0 | 1 | 1 | 30 |
| *Acanthagrion_lancea* | Argentina | Garganta trail | Trp | O | 0 | 0 | 2 | 126 |
| *Acanthagrion_lancea* | Argentina | Yacare Swamp | Trp | O | 0 | 0 | 5 | 70 |
| *Acanthagrion_lancea* | Argentina | Upper Garganta Puddles | Trp | O | 0 | 0 | 6 | 75 |
| *Acanthagrion_minutum* | Guyana | Karanambu dam | Trp | O | 1 | 27 | 42 | 160 |
| *Acanthagrion_trilobatum* | Costa Rica | Natural Lake at Barbilla Lodge | Trp | O | 0 | 1 | 11 | 126 |
| *Acanthagrion_trilobatum* | Costa Rica | Tilapia Pond at Barbilla Lodge | Trp | O | 0 | 0 | 12 | 123 |
| *Aciagrion_bapepe* | Cameroon | Rainforest stream inlet to Dja river | Trp | C | 0 | 1 | 8 | 156 |
| *Aciagrion_nodosum* | Cameroon | Swamp near village after river crossing Dja | Trp | C | 0 | 2 | 1 | 160 |
| *Africallgama_vaginale* | Cameroon | Rainforest trail between village and Dja river | Trp | C | | 0 | 1 | 160 |
| *Africallgama_vaginale* | Cameroon | Swamp near village after river crossing Dja | Trp | C | | 0 | 1 | 75 |
| *Agriocnemis_exilis* | Cameroon | Pond on the route of Banyon | Trp | O | 2 | 6 | 7 | 30 |
| *Agriocnemis_exilis* | Cameroon | Tibati wetland | Trp | O | 2 | 13 | 13 | 60 |
| *Agriocnemis_forcipata* | Cameroon | river at slaughter house | Trp | O | 2 | 7 | 3 | 20 |
| *Agriocnemis_forcipata* | Cameroon | Second stop at Nyong river | Trp | O | 2 | 4 | 6 | 24 |
| *Agriocnemis_victoria* | Cameroon | Second wetland and marsh at Malambe | Trp | O | 2 | 8 | 8 | 20 |
| ... | ... | ... | ... | ... | ... | ... | ... | ... |

### Supporting Methods

#### Female colour classification

Abdominal segments 8-9 are relatively short in damselflies and dragonflies, and males often display brightly-coloured patches in one or both of these segments, which may be absent, duller, or incomplete in females, even when the overall colouration is similar between the sexes (1). In cases where males and females only differed by these last abdomen segments, we coded females as male-coloured. The potential bias of implementing a relatively strict threshold for sexual dimorphism is to reduce the number of sexual dimorphic lineages, which is expected to reduce the rate of transitions from sexual dimorphism to female polymorphism (as there will be fewer sexually dimorphic ancestors). This potential bias runs counter to our hypothesis of how female-limited polymorphism more often evolve (see Introduction), and is therefore conservative.

We note that not in every species has a genetic basis of female polymorphism been confirmed by breeding experiments or molecular studies. Thus, while the data in many cases more accurately describe polychromatisms (i.e. multiple colour types of unknown basis within a population), we here tentatively refer to all female-limited colour variation as female-limited colour polymorphisms. We justify this because in every damselfly species with co-occurring male-coloured and non-male-coloured female types in which discrete colour morphs have been experimentally raised in the laboratory or investigated in depth in the field, a genetic basis of the polymorphism has been confirmed (2–7). We recently characterized the genomic basis of the colour polymorphism that is shared between several species of *Ischnura*, and demonstrated that the alternative alleles underpinning this polymorphism arise from structural variation (i.e. insertions and inversions) at an automosal locus (8). In some species, multiple non-male-coloured female types co-occur without any male-coloured females, but these colour types reflect developmental colour phases, which are often also present in males (9–11). Such purely developmental and non-genetic colour variation was not considered in the present study. For these species, we report only the colour patterns of sexually mature adults. Finally, in no case did we find evidence of male-limited colour polymorphisms.

#### Priors and MCMC simulations

##### Hidden-State Speciation and Extinction

We used a HiSSE model to infer diversifications consequences of sex-related colour states in pond damselflies. We placed equally distributed log-uniform priors (mean = 1e-6, sd = 100) on the three trait-dependent speciation and extinction rates. The effects of the hidden trait on lineage speciation and extinction were drawn from a lognormal with mean in turn sampled from an exponential (rate = 1.0) and standard deviation such that the 95% interval of the hidden rates spans 1 order of magnitude. These hidden rates were then used to scale the trait-dependent diversification rates, resulting in a total of six rates, two (one relatively slow and one relatively fast) for each female-colour state. Transition rates between character states (observed and unobserved) were independently drawn from identical exponential distributions, with mean equal to 10 character transitions across the tree. A birth-death processed conditioned on lineage survival was used as a the tree prior.

We ran two independent replicates of the HiSSE model on the MAP tree from Willink et al. (12). Each replicate was run for 400,000 iterations with a burn-in of 40,000 iterations, with parameter proposals updated every 1,000 iterations. We used stochastic character mapping (13) in these runs to sample character histories and estimate the number of evolutionary origins of female limited polymorphisms in pond damselflies. We also ran a separate HiSSE model without any character state data, to confirm that our results were not driven by the choice of priors. In all phylogenetic analyses, MCMC diagnostics were conducted using the package convenience v 1.0.0 (14) in R v 4.2.2 (15), and ensuring chain convergence and effective sample size (ESS) per run for all model parameters > 300. Ancestral state reconstructions were plotted using the R package RevGadgets v 1.1.0 (16).

##### Bayesian Phylogenetic Mixed Models

We used BPMM to investigate the causal effects of ecological and demographic conditions on the occurrence of female-limited colour polymorphisms. For all multinomial models (*Model 1 - Model 3* and *Model 6*), we used a Kronecker prior (mu = 0, V = units + ) for the fixed effects, as it is relatively flat for the marginal two-way probabilities (i.e. SD vs. SM, SD vs. FP, SM vs. FP) when a logit link function is used (17). We used a parameter-expanded distribution (V = 1, nu = 1,000, alpha.mu = 0, alpha.V = 1) with two degrees of freedom as the prior for the phylogenetic variance components (18). The residual variance, which cannot be identified in multinomial models, was fixed to 1. In all models with field sampled OSR or population density (*Model 3* to *Model 6*), we included the breeding site identity as an additional random effect to account for sampling variance across populations of the same species. We used parameter-expanded priors for this non-phylogenetic random effect (e.g. site; V=1, nu=2, alpha.mu=1, alpha.V=1000). In *Model 5*, we allowed for different population-density slopes in each latitude-habitat combination (tropical-open, tropical-closed, temperate-open, temperate-closed) by fitting the interaction term with sampling effort. For this analysis with a continuous response variable, we assumed an inverse-Wishart prior with a low degree of believe for the residual covariance matrix (V=1, nu=0.002), and we used the default large-variance Gaussian prior (mu = 0, V = I1e+10) for the fixed effects.

For all BPMMs, we ran two independent replicates of each model to diagnose convergence and autocorrelation between posterior samples, using the R package coda (19). We report the posterior means (PM) and 95% highest posterior density intervals (HPD) for parameter estimates. Following Muff et al. (20), we communicate our results in the language of evidence. To ease interpretation, we compute differences in posterior estimates and report the percentage of posterior samples in which the sign of this difference was opposite to our prediction. For example, if we predicted a higher probability of female-limited colour polymorphisms in temperate regions, but female-limited colour polymorphisms occurred with higher probability in tropical regions in 5% of the posterior, we report a PMCMC value of 0.05, as moderate evidence in favour of our prediction.

##### Correlated evolution

We used two separate discrete character models allowing for correlated evolution between latitudinal range and female-colour states and between habitat openness and female-colour states. In both models, transition rates between character states were drawn from independent exponential priors, each with a mean of 10 transitions across the phylogenetic tree. Root-state frequencies were drawn from a Dirichlet prior assuming equal probability of all female-colour and ecological factor combinations. Each model was run in two independent chains, for 400,000 iterations and 40,000 iterations of burn-in, tuning parameter proposals every 1,000. We sampled joint conditional ancestral states and stochastic character maps every 400 iterations. Ancestral state reconstructions in the latitudinal range and female-colour state model strongly supported a most recent common ancestor with monomorphic females occupying a tropical range (see Results). We therefore used the stochastic character maps to estimate the number of event series where a shift to a temperate range predated the origin of female-limited colour polymorphism and the number of event series where the origin of female-limited colour polymorphism predated the temperate shift. We computed the posterior median and 95% HPD interval for each ordered event series, following Landis et al. (21).

##### Relaxed Ornstein-Uhlenbeck

We used a relaxed OU model to infer ancestral shifts in population density optima across the phylogenty of pond damselflies. The stochastic evolution rate ($\sigma^2$) was drawn from a log-uniform distribution bounded between 0.001 and 1. The pull to the optimum ($\alpha$) was drawn from an exponential with mean such that the phylogenetic half-life equals half the length of the tree. The optimum value at the root of the tree ($\theta$) was drawn from a uniform prior bounded between -10 and 10 (i.e. between observing one damselfly every 22,026 survey hours, to observing 22,026 damselflies per survey hour). As we had no prior knowledge of the expected number of density shifts across the tree, we ran three separate analyses with alternative priors of 10, 20, and 40 shifts. When shifts occurred, their size was drawn from a normal distribution with mean of zero and standard deviation such that 95% of shifts were between -1 and 1 in log scale (i.e. between a 63% decrease and 2.7-fold increase in density).

As before, the relaxed OU models were implemented in RevBayes v. 1.1.1 (22), each using two independent chains. MCMC simulations were run for 400,000 iterations and 40,000 iterations of burn-in, tuning parameter proposals every 1,000. We plot the average branch-specific values across the posterior distribution of simulated trees.

### Supporting Results

#### Ecological effects on sex-related colour states

**Table S1**. Posterior mean frequencies of each sex-related colour state in all latitude by habitat openness combinations. Posterior estimates were obtained in *Model 1* (see Methods), using literature data from 297 species of pond damselflies. Abbreviations: SM = sexual monomorphism in colour, FP = female-limited colour polymorphism, SD = sexual dimorphism in colour.

| Latitude | Habitat | SM_mean | FP_mean | SD_mean |
|---|---|---|---|---|
| tropical | open | 0.372 | 0.096 | 0.532 |
| tropical | closed | 0.564 | 0.039 | 0.397 |
| temperate | open | 0.114 | 0.454 | 0.433 |
| temperate | closed | 0.301 | 0.361 | 0.338 |

**Table S2**. Ecological effects on the occurrence of alternative sex-related colour states in pond damselflies. Effects were estimated in *Model 1* (see Methods), using literature data from 297 species. For each state, we contrast latitudinal ranges with similar habitats, and we contrast different habitats at the same latitudinal range. For each comparison we report the posterior mean difference in frequency (PM), the 95% highest posterior density (HPD) interval of differences in frequency, and the percentage of posterior samples in which the difference in frequencies had the opposite sign of the posterior mean (PMCMC). For example, SM occurs with an average of 25.8% (95% HPD interval = 7.6% - 47.0%) higher frequency in tropical-open vs temperate-open habitats. The opposite pattern, of higher frequency of SM in temperate-open habitats was sampled in only 0.04% of the posterior. Abbreviation: SM = sexual monomorphism in colour, FP = female-limited colour polymorphism, SD = sexual dimorphism in colour.

| State | Comparsion | Mean | Lower | Upper | PMCMC |
|---|---|---|---|---|---|
| SM | temperate-open vs tropical-open | -0.2583855 | -0.4700423 | -0.0763370 | 0.004 |
| SM | temperate-closed vs tropical-closed | -0.2628154 | -0.5421785 | 0.0462706 | 0.049 |
| SM | tropical-open vs tropical-closed | -0.1920237 | -0.4099427 | -0.0076624 | 0.033 |
| SM | temperate-open vs temperate-closed | -0.1875938 | -0.4551042 | 0.0344990 | 0.053 |
| FP | temperate-open vs tropical-open | 0.3579481 | 0.1151845 | 0.5922740 | <0.001 |
| FP | temperate-closed vs tropical-closed | 0.3220123 | 0.0601452 | 0.6529775 | 0.001 |
| FP | tropical-open vs tropical-closed | 0.0564585 | -0.0249451 | 0.1656533 | 0.077 |
| FP | temperate-open vs temperate-closed | 0.0923943 | -0.2099826 | 0.4155171 | 0.275 |
| SD | temperate-open vs tropical-open | -0.0995627 | -0.3614189 | 0.1427201 | 0.229 |
| SD | temperate-closed vs tropical-closed | -0.0591969 | -0.3809587 | 0.2892279 | 0.342 |
| SD | tropical-open vs tropical-closed | 0.1355652 | -0.0870159 | 0.3455783 | 0.109 |
| SD | temperate-open vs temperate-closed | 0.0951994 | -0.2365086 | 0.4180145 | 0.277 |

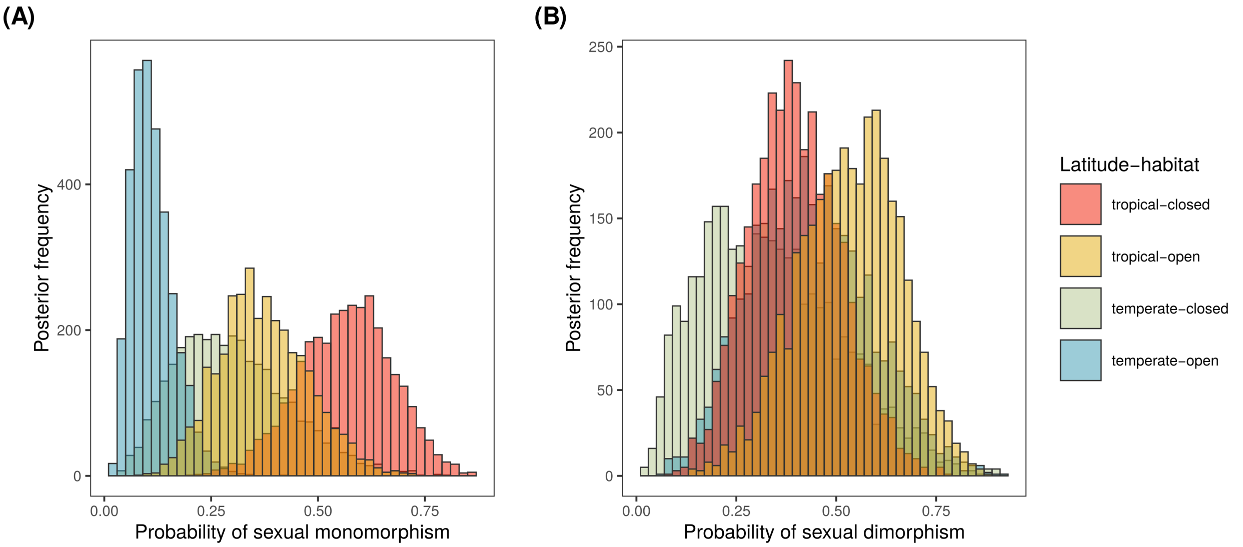

**Figure S1.** Ecological effects on the occurrence of female-monomorphic colour states in pond damselflies. Posterior frequency of **(A)** sexual monomorphism (SM) and **(B)** sexual dimorphism (SD), depending on latitudinal range and habitat openness and based on the Bayesian Phylogenetic Mixed Model (*Model 1*) using literature data. Ecological effects on female-limited polymorphisms are shown in the main text (Fig. 3).

#### Effects of operational sex ration on sex-related colour states

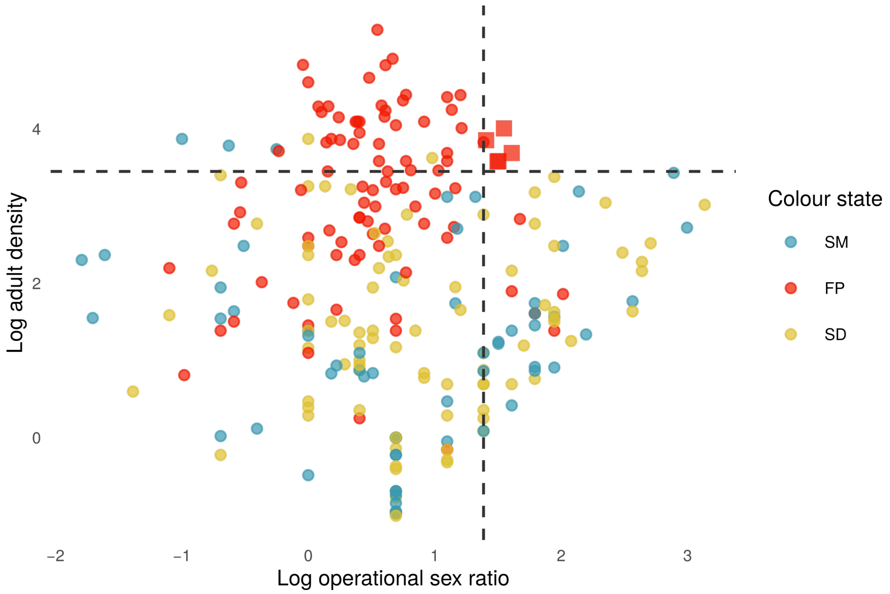

**Figure S2.** Operational sex ratio (OSR) and adult population density in the field data set. Each dot represents a population (n = 208) of a total of 80 species of pond damselflies. Data are log transformed. The dash lines indicate the 80th percentiles for each demographic variable. The square symbols on the top right corner thus correspond to populations in the top 20th percentile for both adult density and male-biased sex ratio. These populations belong to the species *Coenagrion puella*, *Coenagrion pulchellum*, *Enallagma cyathigerum*, and *Xanthocnemis zealandica*. Abbreviation: SM = sexual monomorphism in colour, FP = female-limited colour polymorphism, SD = sexual dimorphism in colour.

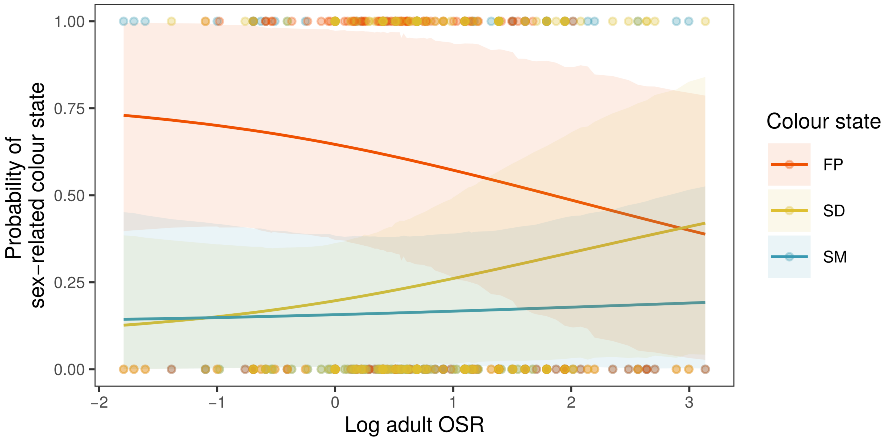

**Figure S3.** Predicted probability of each sex-related colour state with increasing (log) operational sex ratio (OSR; *Model 4*). Larger values indicate more male-biased breeding sites. Shaded areas represent 95% HPD intervals.

#### Ecological effects on demographic mediators

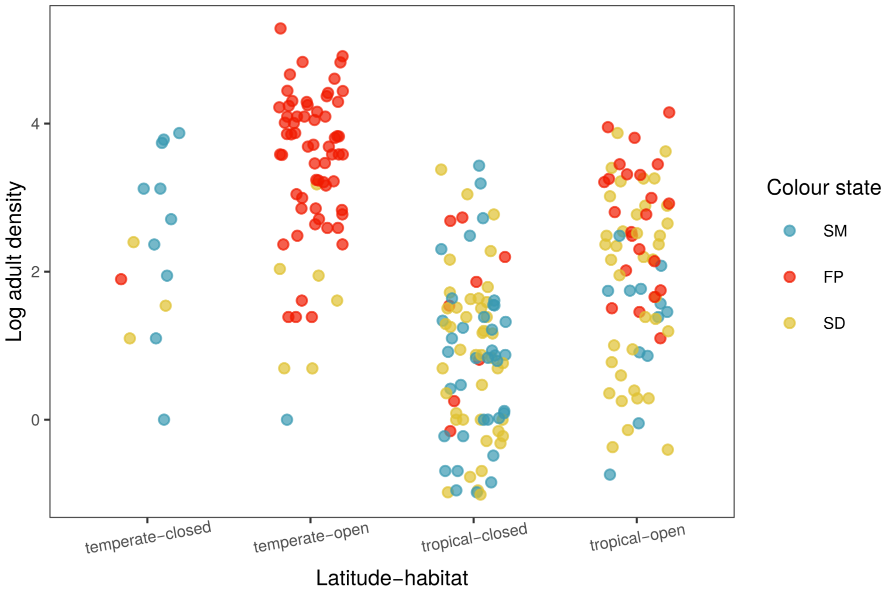

**Figure S4.** Adult population density in all latitude-habitat combinations. Each dot represents a population (n = 208) of a total of 80 species of pond damselflies sampled in the field. Adult density data are log transformed. Abbreviation: SM = sexual monomorphism in colour, FP = female-limited colour polymorphism, SD = sexual dimorphism in colour.

**Table S3**. Ecological effects on adult density at breeding sites in pond damselflies. Effects were estimated in *Model 5* (see Methods), using field data from 208 populations of 80 species. We contrasted latitudinal ranges with similar habitats and different habitats at the same latitudinal range. For each comparison, we report the posterior mean (PM) difference, the 95% highest posterior density (HPD) interval of differences in density, and the percentage of posterior samples in which the difference in density had the opposite sign of the posterior mean (PMCMC). For example, temperate-open habitats have in expectation log(0.977) = 2.63 more individuals than tropical open habitats, for the average sampling effort of 2.6 h. The opposite pattern, of higher adult density in tropical-open habitats, occurred in less than 0.1% of the posterior.

| Comparsion | Mean | Lower | Upper | PMCMC |
|---|---|---|---|---|
| temperate-open vs tropical-open | 0.977 | 0.519 | 1.473 | <0.001 |
| temperate-closed vs tropical-closed | 1.013 | 0.277 | 1.760 | 0.004 |
| tropical-open vs tropical-closed | 0.136 | 0.220 | 1.102 | 0.002 |
| temperate-open vs temperate-closed | 0.644 | -0.011 | 1.276 | 0.023 |

#### Correlated evolution between latitude/habitat occupancy and female-colour states

**Table S4**. Differences in character-state transition rates using a two-trait model allowing for correlated evolution between latitudinal range and female-colour states. We contrasted transitions between female-colour states (FM = female-monomorphic, FP = female-polymorphic) when lineages occupy temperate (Tmp) *vs.* tropical (Trp) ranges. We then contrasted latitudinal shifts between FM and FP lineages. For each rate comparison, we report the posterior median difference, the 95% highest posterior density (HPD) interval of the difference, and the percentage of posterior samples in which the difference was either zero or negative (PMCMC). Note that we here report the posterior median instead of the mean because the correlated evolution model included a reversible jump parameter, switching between independent and correlated evolution. Whenever the MCMC visited the independent model and the sample saved to the posterior, the difference in rates would be exactly zero.

| Transition | Comparison | Median | Lwr | Upr | PMCMC |
|---|---|---|---|---|---|
| FM to FP | Tmp vs Trp | 0.01 | 0.005 | 0.014 | 0.000 |
| FP to FM | Tmp vs trp | 0.00 | 0.000 | 0.006 | 0.640 |
| Tmp to Trp | FM vs FP | 0.00 | -0.001 | 0.004 | 0.713 |
| Trp to Tmp | FM vs FP | 0.00 | 0.000 | 0.004 | 0.703 |

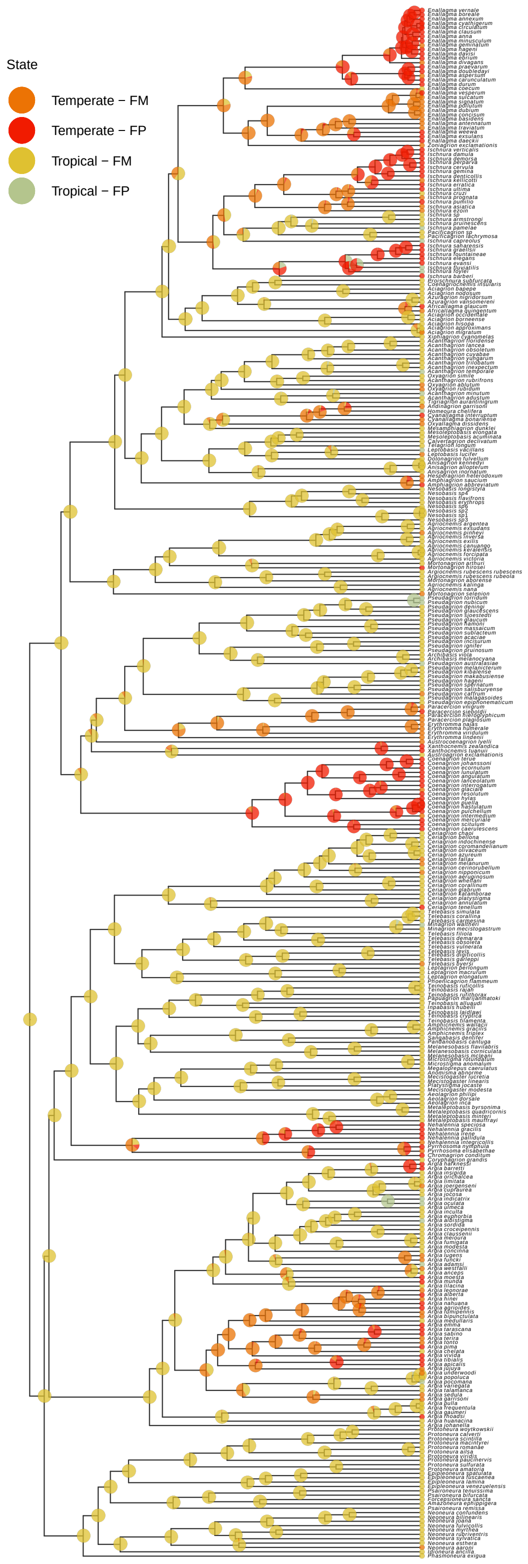

**Figure S5.** Ancestral state reconstruction of latitudinal ranges and female colour states in pond damselflies (family Coenagrionidae). Colours at the tips show colour states in extant taxa. Pies indicate indicate joint conditional posterior probabilities of female colour states in ancestral nodes. Abbreviations: FM = female monomorphism, FP = female-limited colour polymorphism.

**Table S5**. Differences in character-state transition rates using a two-trait model allowing for correlated evolution between habitat openness and female-colour states. We contrasted transitions between female-colour states (FM = female-monomorphic, FP = female-polymorphic) when lineages occupy open (HO) *vs.* closed (HC) ranges. We then contrasted habitat shifts between FM and FP lineages. For each rate comparison, we report the posterior median difference, the 95% highest posterior density (HPD) interval of the difference, and the percentage of posterior samples in which the difference was either zero or negative (PMCMC). Note that we here report the posterior median instead of the mean because the correlated evolution model included a reversible jump parameter, switching between independent and correlated evolution. Whenever the MCMC visited the independent model and the sample saved to the posterior, the difference in rates would be exactly zero.

| Transition | Comparison | Median | Lwr | Upr | PMCMC |
|---|---|---|---|---|---|
| FM to FP | HO vs HC | 0.005 | 0.003 | 0.007 | 0.000 |
| FP to FM | HO vs HC | 0.000 | 0.000 | 0.005 | 0.909 |
| HC to HO | FM vs FP | 0.000 | -0.001 | 0.003 | 0.640 |
| HO to HC | FM vs FP | 0.000 | 0.000 | 0.000 | 0.989 |

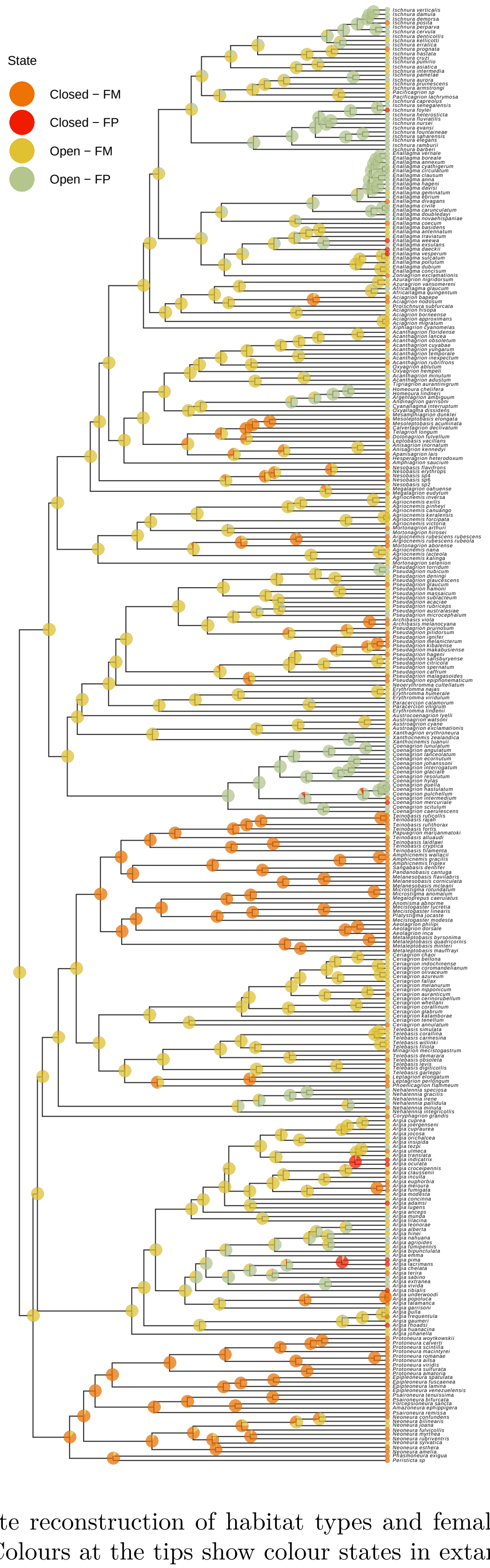

**Figure S6.** Ancestral state reconstruction of habitat types and female colour states in pond damselflies (family Coenagrionidae). Colours at the tips show colour states in extant taxa. Pies indicate indicate joint conditional posterior probabilities of female colour states in ancestral nodes. Abbreviations: FM = female monomorphism, FP = female-limited colour polymorphism.

#### Evolution of adult density optima

We estimated adult density optima using relaxed Ornstein-Uhlenbeck models. These models were parametrized with the prior expected number of shifts in adult density across the pond-damselfly tree (see methods). Since we had no prior knowledge on the expected number of shifts, we ran separate models with three alternative priors: 10, 20, and 40 shifts. Results for the 20-shift prior are shown in the main text, and results for the other two models are shown below (Fig. S7-S8). In all models, adult densities were inferred to increase after the common ancestor of clades which are currently dominated by female-polymorphic taxa (Table S8). Namely, adult densities increased after the common ancestor of the clade including *Nehalennia*, *Pyrrhosoma* and *Chromagrion*, the genus *Coenagrion*, the genus *Xanthocnemis*, the clade including *Homeoura* and *Argentagrion*, the clade including *Ischnura* and *Enallagma*, and then again in the genus *Ischnura* (Fig. 4d; Fig. S7-S8). More subtle density shifts were noted on the branches leading to *Acanthagrion minutum* and its nearest relative, *Leptobasis vacillans*, and *Pseudagrion nubicum*, especially in models with a higher *a priori* total number of transitions (Fig. 4d; S7-S8). These are three FP species within clades that are majority FM (Table S8).

The genus *Argia* is as an exception to this general trend, as adult densities in the genus are largely conserved and intermediate (Fig. 4d; S7-S8). Two other exceptions, *Agriocnemis* and *Erythromma*, are female-monomorphic clades descending from branches with positive shifts in density. Interestingly, *Agriocnemis* females exhibit developmental colour phases that reduce male mating harassment prior to reproductive maturation (9), and *Erythromma* has both male-coloured and non-male-coloured female forms, but in this study, we conservatively coded them as female-monomorphic because the literature is explicitly ambiguous as to whether these forms are developmental phases or genetic morphs (23).

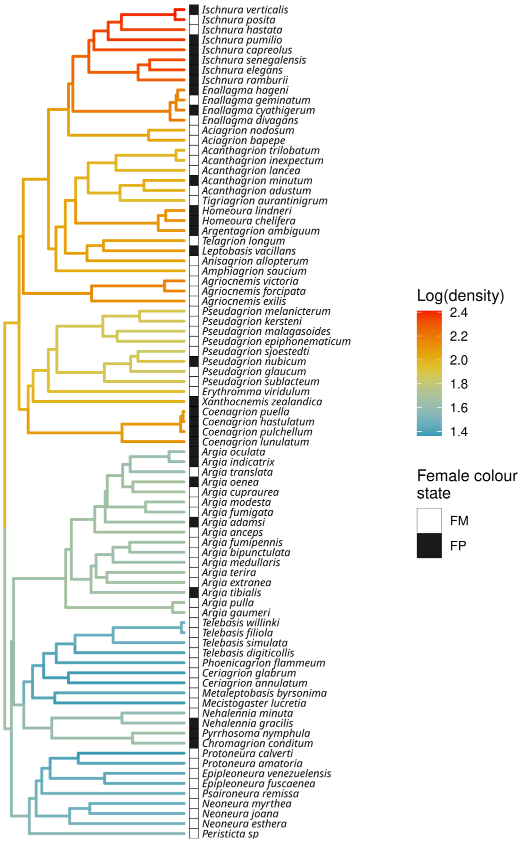

**Figure S7.** Evolution of adult density optima ($\theta$) based on a relaxed Ornstein-Uhlenbeck model. The model was parametrized with a prior expectation of **10** optima shifts across the tree. The phylogeny includes only the taxa sampled in the field (n = 83). Multiple density surveys were average and density was log-transformed before the analysis.

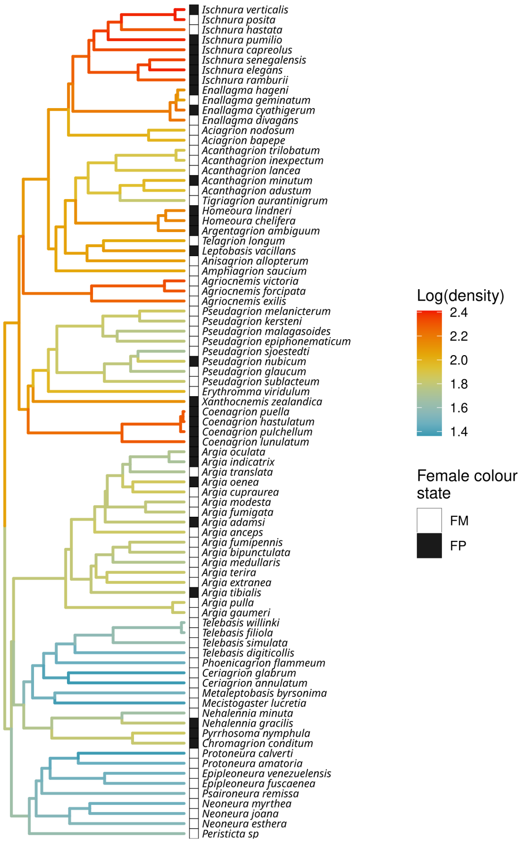

**Figure S8.** Evolution of adult density optima ($\theta$) based on a relaxed Ornstein-Uhlenbeck model. The model was parametrized with a prior expectation of **40** optima shifts across the tree. The phylogeny includes only the taxa sampled in the field (n = 83). Multiple density surveys were average and density was log-transformed before the analysis.

**Table S6**. Pond damselfly clades with branch shifts in adult density optima, based relaxed OU models. For each clade with FP taxa and evidence of branch shifts in adult density optima (Fig. 4d; S7-S8), we show whether the shift directly followed the origin of the clade (MRCA) or if it was inferred in a shallower branch (tip branch). We also present the total number of species for which we obtained female-colour data, and the number and proportion of those species that are FP. The family wide proportion of FP taxa is 0.28. Finally, we provide the total number of species in the clade following the *World Odonata List* curated at the Puget Sound Museum of Natural History. We include the genus *Oxyagrion* within *Acanthagrion*, as *Acanthagrion* is likely otherwise paraphyletic (12), and *Amorphostigma* and *Pacificagrion* within *Ischnura* (24).

| Clade | Shift location | Spp with data | FP spp | FP proportion | Total spp |
|---|---|---|---|---|---|
| *Nehalennia + Pyrrhosoma + Chromagrion* | MRCA | 9 | 6 | 0.67 | 9 |
| *Coenagrion* | MRCA | 17 | 15 | 0.88 | 28 |
| *Xanthocnemis* | MRCA | 2 | 2 | 1.00 | 2 |
| *Homeoura + Argentagrion* | MRCA | 3 | 3 | 1.00 | 5 |
| *Enallagma + Ischnura* | MRCA | 71 | 45 | 0.63 | 120 |
| *Ischnura* | MRCA | 37 | 24 | 0.65 | 76 |
| *Pseudagrion* | tip branch | 33 | 3 | 0.09 | 165 |
| *Leptobasis + Mesoleptobasis + Calvertagrion + Dolonagrion + Tealagrion* | tip branch | 8 | 1 | 0.12 | 21 |

*(continued)*

| Clade | Shift location | Spp with data | FP spp | FP proportion | Total spp |
|---|---|---|---|---|---|
| *Acanthagrion* | tip branch | 17 | 2 | 0.12 | 68 |

#### Background heterogeneity in diversification

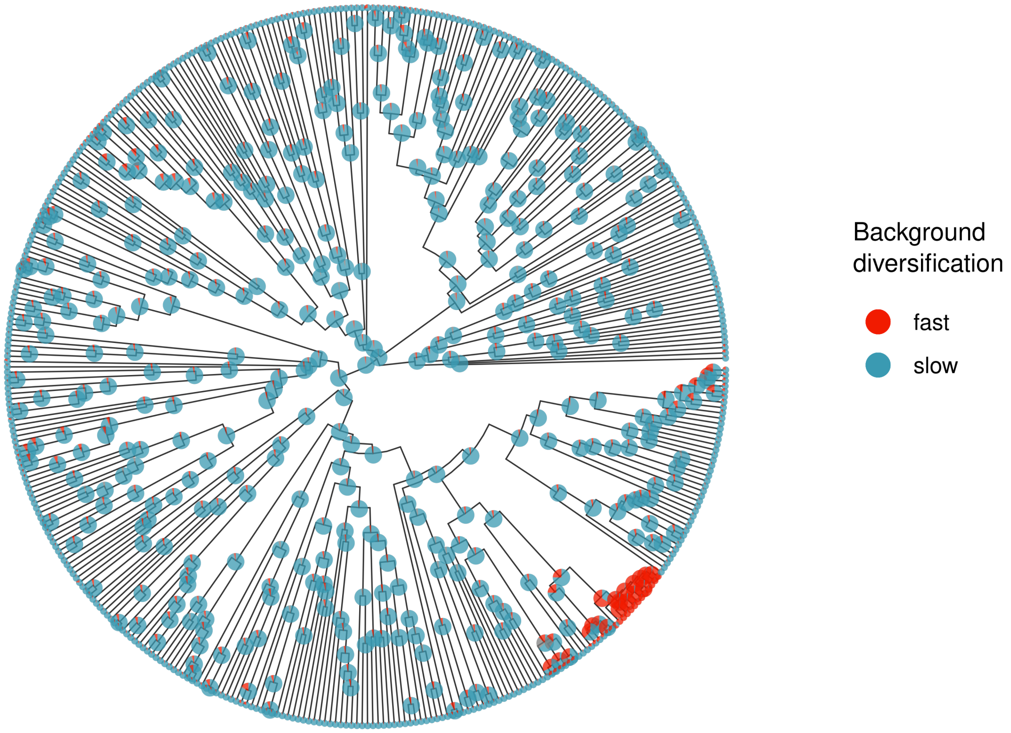

**Figure S9.** Ancestral state estimation (ASE) for a hidden trait controlling background heterogeneity in diversification. ASE is base on a Hidden-State Speciation and Extinction (HiSSE) model. The hidden trait include two states, for slow and fast background diversification respectively. Fast background diversification is inferred primarily in the North American Bluets (genus *Enallagma*) and, to a lesser extent, in a clade of North American Forktails (genus *Ischnura*).

#### Diversification dynamics

**Table S7**. Differences in diversification rates between alternative sex-related colour states, using a Hidden-State Speciation and Extinction (HiSSE) model. For each rate type, we contrasted sexually monomorphic

(SM), female polymorphic (FP), and sexually dimorphic (SD) lineages, with either low (0) or high (1) background diversification rates ("Hidden"). For each rate comparison, we report the posterior mean difference (PM), the 95% highest posterior density (HPD) interval of the difference, and the percentage of posterior samples in which the difference had the opposite sign of the posterior mean (PMCMC). For example, the difference in speciation rates between FP and SM lineages with a low background rate of diversification is 0.065 (95% HPD interval = 0.021 - 0.116). The opposite pattern, of higher speciation in SM lineages occurred in less than 0.1% of the posterior.

| Rate_type | Comparison | Hidden | Mean | Lwr | Upr | PMCMC |
|---|---|---|---|---|---|---|
| speciation | SM vs FP | 0 | -0.065 | -0.116 | -0.021 | <0.001 |
| speciation | SM vs FP | 1 | -0.267 | -0.486 | -0.068 | <0.001 |
| speciation | SM vs SD | 0 | -0.020 | -0.036 | -0.004 | 0.004 |
| speciation | SM vs SD | 1 | -0.090 | -0.191 | -0.005 | 0.004 |
| speciation | SD vs FP | 0 | -0.045 | -0.092 | -0.001 | 0.009 |
| speciation | SD vs FP | 1 | -0.177 | -0.375 | -0.008 | 0.009 |
| extinction | SM vs FP | 0 | -0.045 | -0.102 | 0.006 | 0.064 |
| extinction | SM vs FP | 1 | -0.094 | -0.263 | 0.025 | 0.064 |
| extinction | SM vs SD | 0 | 0.000 | -0.013 | 0.010 | 0.482 |
| extinction | SM vs SD | 1 | -0.004 | -0.030 | 0.026 | 0.482 |
| extinction | SD vs FP | 0 | -0.045 | -0.104 | 0.007 | 0.076 |
| extinction | SD vs FP | 1 | -0.090 | -0.291 | 0.022 | 0.076 |
| net diversification | SM vs FP | 0 | -0.019 | -0.048 | 0.007 | 0.067 |
| net diversification | SM vs FP | 1 | -0.173 | -0.399 | 0.014 | 0.04 |
| net diversification | SM vs SD | 0 | -0.020 | -0.035 | -0.005 | 0.004 |
| net diversification | SM vs SD | 1 | -0.087 | -0.194 | -0.006 | 0.011 |
| net diversification | SD vs FP | 0 | 0.001 | -0.027 | 0.028 | 0.512 |
| net diversification | SD vs FP | 1 | -0.086 | -0.286 | 0.074 | 0.151 |

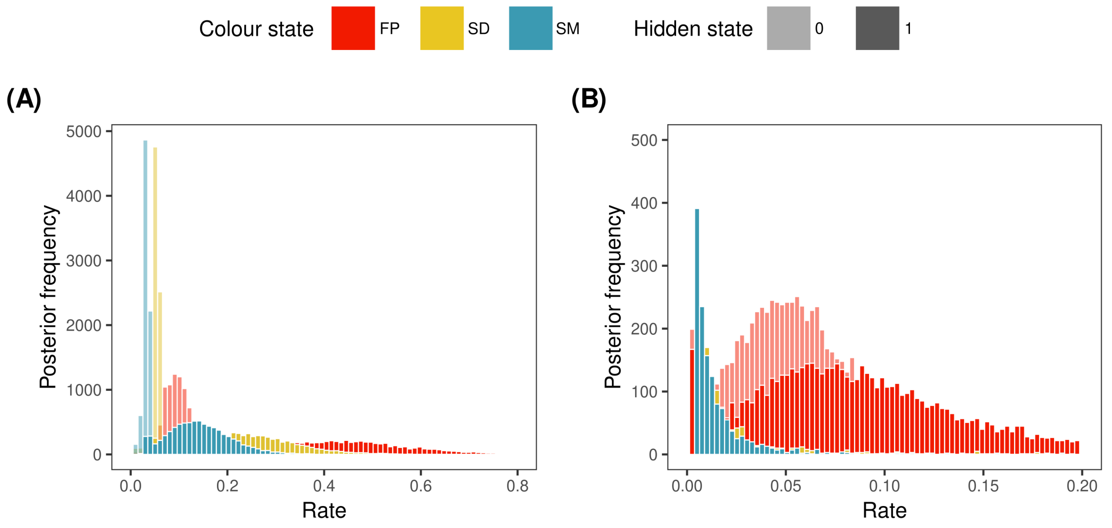

**Figure S10.** State-dependent speciation and extinction rates in pond damslelflies. Histograms represent posterior distributions from a Hidden State-dependent Speciation and Extinction (HiSSE) model. Background heterogeneity in diversification was modeled as a hidden trait with two values (0 = low background diversification, 1 = high background diversification). **(A)** Speciation. **(B)** Extinction. Abbreviations: FP = female-limited polymorphism, SD = sexual dimorphism, SM =sexual monomorphism.

#### Evolutionary transitions between sexual monomorphism and sexual dimorphism

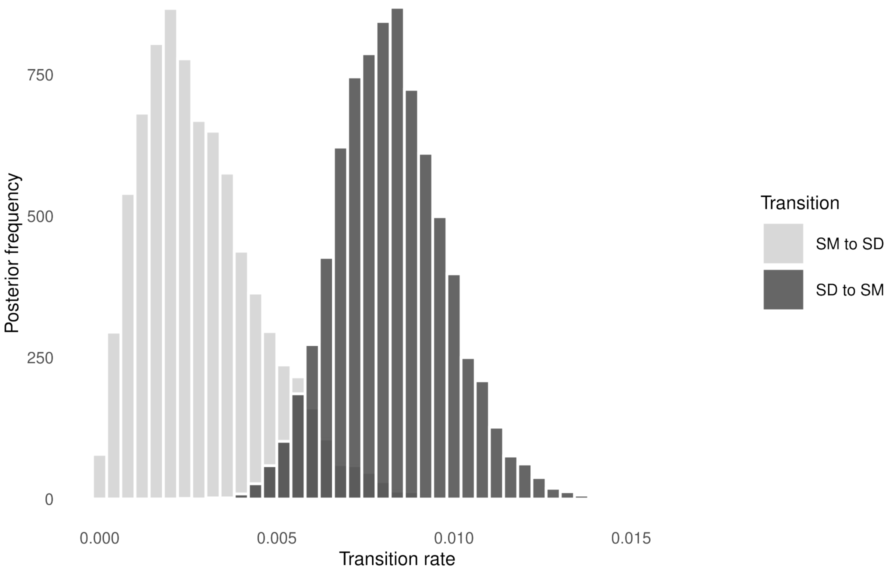

**Figure S11.** Transition rates between sexual monomorphism in colour (SM) and sexual dimorphism in colour (SD) in pond damselflies. The histograms show posterior distributions based on a Hidden State Speciation and Extinction (HiSSE) model.
